# Transcriptome and proteome associated to the sex-dependent effects of Neuroligin-2 absence on wakefulness and sleep features

**DOI:** 10.1101/2025.07.25.666775

**Authors:** Nicolas Lemmetti, Tanya Leduc, Julien Dufort-Gervais, Jean-Marc Lina, Valérie Mongrain

## Abstract

Neuroligin-2 (NLGN2) regulates GABAergic neurotransmission and is linked to neurodevelopmental disorders. The absence of NLGN2 in male mice decreases time spent in slow wave sleep and alters electrocorticographic (ECoG) activity. We tested whether NLGN2 absence also impacts wake/sleep states in females and whether sleep phenotypes associate with specific omic signatures of the cerebral cortex. *Nlgn2* knockout (KO) mice and wild-type (WT) littermates were implanted with ECoG electrodes, and ECoG signals were recorded for 48 hours. Other cohorts were submitted to motor cortex sampling followed by quantifications of the transcriptome or proteome. KO mice of both sexes spent less time in slow wave and paradoxical sleep, and KO males (but not females) showed more wake and slow wave sleep episodes during the light period. Mutant animals of both sexes showed widespread differences in wake/sleep ECoG spectral activity when compared to WT mice, including a slower theta peak frequency during paradoxical sleep. The most prominent Hurst exponent was significantly increased in *Nlgn2* KO animals during all states, and Hurst exponents were less dispersed during paradoxical sleep only in KO females. Less than 31% of genes with expression changed in KO versus WT mice were shared between females and males (e.g., linked to MAPK/ERK pathway, protein regulation and neurotransmission), and it was less than 15% in the case of proteins (e.g., related to calcium signaling and excitatory synapse function). Moreover, KO males showed a higher number (> 2.5 fold) of genes with expression significantly increased and decreased compared to control mice than KO females, but only KO females presented significantly lower levels of a subset of core proteins regulating synapses (e.g., SYNGAP1, HOMER1). The findings indicate that effects of NLGN2 absence on wake/sleep phenotypes and omic landscapes depend on biological sex, and could help understanding the origin of sleep disturbances in neurodevelopmental disorders.

## 1 Introduction

Neuroligins (NLGNs) contribute to postsynaptic density specialization via interactions with scaffolding proteins and receptors (Barrow et al., 2009; Budreck et al., 2013; Poulopoulos et al., 2009; Sui et al., 2024). Their roles in neurotransmission and stabilization of pre- and postsynaptic terminals depend on the binding with presynaptic partners, in particular neurexins (Tsetsenis et al., 2014). Mutations (and dysregulations) of NLGNs have been linked to neurodevelopmental disorders, such as autism spectrum disorder (ASD) and schizophrenia (Jamain et al., 2003; Nakanishi et al., 2017; Sun et al., 2011), and to neurological conditions like Alzheimer’s disease (Arias-Aragón et al., 2023). All these brain diseases associate with sleep disturbances (Freeman et al., 2020; Schwichtenberg et al., 2022; Zhang et al., 2022), and have a prevalence highly dependent on biological sex. Although modifications in wake/sleep architecture and electrocorticographic (ECoG) activity characterize animals lacking individual NLGNs (Areal et al., 2025; Cao et al., 2020; El Helou et al., 2013; Leduc et al., 2024; Li et al., 2013; Liu et al., 2017; Massart et al., 2014; Seok et al., 2018; Thomas et al., 2017), the impact of sex on this relationship remains mostly undefined.

NLGN2 is predominantly found at inhibitory (γ-aminobutyric acid (GABA)-ergic) synapses (Varoqueaux et al., 2004; Chih et al., 2006). It contributes to the recruitment and clustering of gephyrin and GABA_A_ receptors at the postsynaptic density (Dong et al., 2007; Poulopoulos et al., 2009), and to the normal function and maintenance of inhibitory synapses (Chen et al., 2020; Groisman et al., 2023; Lech et al., 2026; Liang et al., 2015; Troyano-Rodriguez et al., 2019). The absence of NLGN2 in male mice was found to, among others, decrease the time spent in slow wave sleep (SWS), increase the density and amplitude of sleep slow waves (SW), and accelerate paradoxical sleep (PS) recovery following sleep deprivation (SD) (Leduc et al., 2024; Seok et al., 2018). Moreover, aperiodic properties of the wake and sleep ECoG signal are also altered in *Nlgn2* knockout (KO) male mice that notably show higher long-range dependency for all states (Leduc et al., under review). ECoG aperiodic analyses complement standard spectral analysis interrogating brain oscillations and further inform on brain information processing since aperiodic metrics can predict changes in working memory and cognitive scores in patients with neurodegenerative diseases (Averna et al., 2023; Donoghue et al., 2020). Importantly, whether the wide range of wake and sleep ECoG phenotypes found in *Nlgn2* knockout (KO) mice associates with specific signatures of the cerebral cortex transcriptome and proteome is an open question, to which answering will help understanding the underlying mechanisms.

Sex differences in sleep phenotypes have been described in healthy human subjects and rodents (Carrier et al., 2017; Choi et al., 2021; Dib et al., 2021; Ehlen et al., 2013; Koehl et al., 2006; Mongrain et al., 2005; Paul et al., 2006; Swift et al., 2020; van den Berg et al., 2009). In addition, patients affected with multiple neurodevelopmental disorders express a symptomatology that varies with sex, including when considering sleep phenotypes (Bricout et al., 2024; Estes et al., 2023; Hartley & Sikora, 2009; Robison et al., 2008; Saré & Smith, 2020; Weir et al., 2021). NLGNs are encoded by five different genes in humans with *NLGN3* to *5* that are located on sex chromosomes (Bolliger et al., 2001; Kikuno et al., 1999; Nguyen et al., 2020; Philibert et al., 2000; Skaletsky et al., 2003), and by four genes in rodents with *Nlgn3* and *Nlgn4* being on sex chromosomes (Bolliger et al., 2008; Maxeiner et al., 2020). Also, sex hormones impact the expression of *Nlgn* genes and NLGN protein levels in neurons (Bethea & Reddy, 2012; Sellers et al., 2015; Srancikova et al., 2021; Ziehn et al., 2012). Studies of mice lacking NLGN1 suggest that the effects on sleep phenotypes may depend on sex (Areal et al., 2025; El Helou et al., 2013; Massart et al., 2014). However, whether the absence of NLGN2 impacts wake/sleep states differently in male and female mice remains to be established, together with the corresponding omic landscapes.

The hypothesis tested in this study was that the absence of NLGN2 in mice impacts wake/sleep architecture and ECoG spectral and aperiodic metrics differently in females and males, which is coupled to sex-specific transcriptomic and proteomic signatures in mutant animals. This was investigated by recording the motor and visual cortex ECoG signal of male and female *Nlgn2* KO mice and wild-type (WT) littermates during undisturbed conditions and following SD, to investigate the effects of a sleep homeostatic challenge, and by measuring genome-wide gene expression and protein abundance of the motor cortex. The architecture of wake/sleep states (including state alternation) was found to be differently altered in male and female *Nlgn2* KO mice. Moreover, *Nlgn2* KO mice of both sexes showed differences in ECoG spectral and multifractal activities, with KO females being generally more affected than KO males when considering PS. The transcriptome was observed to be particularly impacted by the mutation in KO males (i.e., higher number of genes with expression significantly increased and decreased in males than females), whereas an ensemble of key proteins with roles at excitatory synapses appear notably affected only in KO females (e.g., SHANK3, HOMER1, GLUA3 level significantly lower in KO females but not in KO males). This study highlights that the effects of NLGN2 absence on wake/sleep phenotypes and associated omic signatures differ between sexes. Given the association of NLGN2 with neuropsychiatric diseases, these findings could help to understand the mechanisms underlying sleep disturbances in neurodevelopmental disorders such as ASD.

## 2 Methods

### 2.1 Animals

*Nlgn2* mutant mice were obtained from The Jackson Laboratory (Bar Harbor, ME, USA; B6;129-*Nlgn2*^tm1Bros^/J; RRID: IMSR_JAX:008139) and backcrossed for more than 10 generations to C57BL/6J mice (Jackson Laboratory; RRID: IMSR_JAX:000664). The mutation results in an absence of NLGN2 protein due to the deletion of the first coding exon, as described previously (Varoqueaux et al., 2006). Mice heterozygous for the mutation were bred to obtain WT, heterozygous and *Nlgn2* KO homozygous mice. Male and female pups were separated at weaning and housed in groups of two to five littermates per cage (access to food and water *ad libitum*). Animals were maintained under a 12-hour light/12-hour dark cycle and between 22-25 °C throughout breeding and experimental procedures. For electrophysiology, mice were individually housed starting at least two weeks before surgery and the following groups have been studied: eleven WT females (age and weight at surgery: 79.2 ± 6.5 days, 20.6 ± 1.3 g), nine WT males (78.9 ± 3.0 days, 26.7 ± 1.2 g), ten KO females (80.4 ± 4.5 days, 19.2 ± 0.8 g), and ten KO males (77.5 ± 4.7 days, 22.5 ± 1.7 g). The age at surgery was not significantly different between groups (Genotype-by-Sex interaction F_1,36_ = 0.4, p = 0.5; main Genotype effect F_1,36_ = 0.002, p = 1.0; main Sex effect F_1,36_ = 0.6, p = 0.4). However, the weight at surgery was significantly lower in KO males compared to WT males (Genotype-by-Sex interaction F_1,36_ = 5.3, p = 0.03; p < 0.0001 between KO and WT males; p = 0.09 between KO and WT females), as we have reported previously for a different cohort of *Nlgn2* KO male mice having a mixed genetic background (B6;129; Leduc et al., 2024; Seok et al., 2018). For the transcriptome quantification, four WT females, three KO females, four WT males and three KO males were used (all between 74 and 98 days at sacrifice), while it was four WT females, four KO females, five WT males, and three KO males (all between 63 and 112 days) for the quantification of the proteome. Sample sizes were defined according to previously published datasets using similar experimental designs, outcome variables and/or animal model (Areal et al., 2025; Franken et al., 2006; Leduc et al., 2024; Noya et al., 2019; Paul et al., 2006; Seok et al., 2018).

### 2.2 Experimental protocols

For electrophysiology, mice were implanted with electrodes as detailed below, and then recovered for about five days and habituated to cabling conditions for at least seven days before recordings. ECoG and electromyographic (EMG) recordings started at light onset (defined as Zeitgeber time 0: ZT0) and lasted for 48 hours including 24 hours of undisturbed conditions (referred to as the baseline day: BL), followed by a 6-hour SD (starting at ZT0), and 18 hours of undisturbed conditions (SD together with following recovery referred to as the recovery day: REC). SD was achieved by delicately approaching a serological pipette near the mouse or by moving the litter next to the mouse only when the animal was showing signs of sleep (e.g., immobility, sleep posture, etc.) to minimize the stress experienced by the animals. SD also involved moving the mouse to a cage with clean litter at the beginning of the fourth hour, and adding paper tissue to the cage during the third and sixth hours only if sleep was increasingly difficult to prevent. For omic quantifications, all mice were sacrificed under undisturbed conditions between ZT1 and ZT3 (i.e., between 1 and 3 hours after light onset) using a random order of sacrifice. Brains were extracted following guillotine decapitation without anesthesia, and motor cortices were rapidly dissected on ice. Motor cortices were immediately frozen on dry ice for subsequent RNA quantification or in liquid nitrogen for protein quantification. Experimental procedures were conducted according to the guidelines of the Canadian Council on Animal Care and were approved by the *Comité d’éthique de l’expérimentation animale du Centre intégré universitaire de santé et services sociaux du Nord-de-l’Île-de-Montréal* or by the *Comité institutionnel de protection des animaux* of the *Centre de recherche du Centre hospitalier de l’Université de Montréal*.

### 2.3 Electrode implantation surgery

*Nlgn2* KO mice and WT littermates were implanted with ECoG and EMG electrodes as described previously (Areal et al., 2020; Dufort-Gervais et al., 2021; Freyburger et al., 2016). This was done in batches of four to eight animals, always including both genotypes instrumented in a random order. Briefly, animals were anesthetized with a mix of Ketamine/Xylazine (120/10 mg/kg, i.p. injection) prior to the surgery, and stable anesthesia was maintained with isoflurane (0.5 to 1% in 2 L/min O_2_). ECoG electrodes consisted of two gold-plated screws (1.14 mm thread major diameter) that were screwed through the skull over the right cerebral hemisphere (motor cortex: 1.5 mm lateral to midline, 1.5 mm anterior to bregma; visual cortex: 1.5 mm lateral to midline, 1.0 mm anterior to lambda). An additional screw was positioned on the right hemisphere and served as a reference electrode (2.6 mm lateral to midline, 0.7 mm posterior to bregma). Over the left hemisphere, three anchor screws were implanted to maximize adherence of the montage to the skull. Additionally, two gold wires (0.2 mm diameter) were inserted in neck muscles to serve as EMG electrodes. The ECoG and EMG electrodes were soldered to a connector and fixed to the skull by cementing the montage (including anchor screws) with dental cement. Detailed information about material used for electrode implantation can be found elsewhere (Dufort-Gervais et al., 2021).

### 2.4 ECoG recording and wake/sleep architecture analysis

ECoG/EMG signals were recorded, amplified (Lamont amplifiers), sampled at 256 Hz, and filtered using the commercial software Harmonie (Natus, Middleton, WI, USA), as done previously (Areal et al., 2020; Freyburger et al., 2016; Seok et al., 2018). Bipolar signals were segmented into 4-second epochs and visually inspected to assign a state of vigilance (wakefulness, SWS and PS) to each epoch based on ECoG and EMG characteristics as detailed previously (Areal et al., 2025; Franken et al., 1999; Mang & Franken, 2012). Visual assignment of vigilance states was done blind to genotype. In brief, wakefulness was identified by an ECoG signal of relatively low amplitude (in comparison to SWS) and with activity in a broad range of frequencies, along with a variable EMG signal generally of high amplitude. SWS was characterized by an ECoG signal of high amplitude and a low amplitude EMG signal. PS was defined by an ECoG signal dominated by regular theta waves (6-9 Hz) and by an EMG signal showing the lowest amplitude (relative to wake and SWS). Epochs containing artifacts were identified simultaneously and later excluded from ECoG activity analyses. The time spent in vigilance state was calculated per hour and for the 12-hour light and 12-hour dark periods for both BL and REC. The number of episodes of each state and the mean duration of individual episodes of vigilance states were averaged for the 12-hour light and 12-hour dark periods for BL and REC. For REC, the latency to SWS following SD was computed as the time elapsed from the end of the SD to the first 60-second episode of SWS that was not interrupted by more than two non-consecutive 4-second epochs of wakefulness. The latency to PS was defined as the elapsed time between the end of the SD and the first 4-second epoch of PS as previously done (Vassalli & Franken, 2017). The latency difference was calculated as the difference between PS latency and SWS latency. Finally, the effect of SD and recovery on SWS and PS duration (rebound) was assessed using accumulated differences from the corresponding BL values. The slope of recovery was calculated for SWS and PS accumulated differences between ZT6 (i.e. the end of SD after which recovery begins) and ZT12 (i.e. the end of the light period which is the usual rest period for nocturnal animals), as done previously (Leduc et al., 2024).

### 2.5 Spectral analysis of the ECoG

Spectral power between 0.75-50 Hz was computed by applying a Fast Fourier transform (with mean subtraction and extended cosine bell) on the bipolar ECoG signal (motor - visual) of artifact-free 4-second epochs (4-second windows aligned to state epochs and no overlap between windows). The same was also done for signals from individual electrodes to obtain power spectra specific to the electrodes positioned above the motor and visual cortices (i.e., motor - reference and visual - reference). Absolute spectral power was compared between groups. To account for interindividual differences in overall ECoG power, ECoG spectra were also expressed relatively to the mean total ECoG power of all frequencies of all states during BL for each mouse. Only epochs visually identified as quiet wake (low amplitude EMG) were used for wakefulness spectral analysis to avoid contamination of the ECoG signal by EMG activity. Spectral activity was also computed separately for delta activity (1-4 Hz) during SWS, and theta (6-9 Hz) and alpha (9-12 Hz) frequency bands during wakefulness. To align spectral data to the distribution of wake and sleep for each animal, the power in these spectral bands was averaged per interval each comprising an equal number of wake or SWS epochs as done previously (Areal et al., 2020; Freyburger et al., 2016). More precisely, for SWS during BL, delta power was calculated for 12 and six intervals during the light and dark periods, respectively. During REC, SWS delta power was calculated for eight intervals during the 6-hour light period after the end of SD and for six intervals during the dark period. Given that artifact-free epochs are less frequent for wakefulness given the high EMG tone, spectral power was computed for three and six intervals during the light and dark periods of BL, respectively. For REC, spectral power in wake was computed for four intervals during the SD, one interval during the remaining 6 hours of the light period after the end of SD, and six intervals during the dark period. The 24-hour dynamics was calculated for relative power in spectral bands, which was computed by expressing the power per interval of BL and REC as a percentage of the mean 24-hour BL spectral power in the targeted band for each animal.

### 2.6 Detection and analyses of slow waves

Detection of individual SW was done on the bipolar ECoG signal of SWS artifact-free epochs after band-pass filtering (between 0.5 and 4.0 Hz) using a linear phase FIR filter of −3 dB (Freyburger et al., 2017; Leduc et al., 2024; Massart et al., 2014). Automatic detection of SW was performed using the following criteria: negative-to-positive peak-to-peak amplitude > 100 µV, negative peak amplitude > 40 µV, negative phase duration between 0.1 and 1.0 second and positive phase duration < 1.0 second. The criteria for automatic detection are the same as those used recently, which were specifically optimized for detection in *Nlgn2* KO mice (Leduc et al., 2024). SW density was calculated as the number of SW per minute of SWS. For each SW, four other metrics were derived: peak-to-peak amplitude (voltage difference between the negative and positive peaks of unfiltered signal [µV]), slope (velocity of the change between the negative and positive peaks [µV/second]), duration of the negative phase (second), duration of the positive phase (second). SW metrics were computed for the 24 hours of BL and REC. To interrogate their 24-hour dynamics, these metrics were also averaged for the same intervals described above for SWS delta activity.

### 2.7 Multifractal analysis of the ECoG

Scaling properties and regularities/irregularities across time scales of the ECoG were evaluated using a multifractal analysis similar to previously conducted (Areal et al., 2025; Hector et al., 2026; Leduc et al., under review; Lina et al., 2019). The multifractal framework (based on statistics of multiscale fluctuations) reveals local scale-invariant properties in the ECoG signal. It extends the spectral analysis of 1/f^γ^ processes by characterizing the distribution of the instantaneous spectral decay exponents (here for the frequency range 0.5 to 64 Hz). Using a Wavelet-Leaders formalism developed previously (Ciuciu et al., 2008; Wendt et al., 2009), the Hurst exponent (*H*) was obtained from the scaling exponent γ = 2*H* + 1 and using a Daubechies wavelet with six vanishing moments. For each artifact-free 4-second epoch, two parameters were extracted: the most prevalent *H* (*Hm*) and the *Dispersion* of all *H* around *Hm* (i.e., the range of all *H* values for a given epoch) (Jaffard et al., 2006; Lina et al., 2019). *Hm* informs about long-range dependency (temporal correlation) of the power across time scales of the ECoG signal, with lower values of *Hm* indicating more anti-persistence (less long-range dependency) and higher values less anti-persistence (higher long-range dependency) (Areal et al., 2025; Hector et al., 2026; Lina et al., 2019). *Dispersion* describes the *Hm* variability when considering local points in time, and was used as an indicator of “local instability” (or “multifractality”). *Dispersion* closer to 0 is interpreted as higher stability (of the power decay with exponent γ across frequencies) since it is driven by less variability of *Hm* at a given timepoint. Conversely, more negative *Dispersion* is indicative of lower local stability because it originates from more variability of *Hm*. The two metrics were computed for the full 24-hour BL and 24-hour REC, and their 24-hour dynamics were also interrogated using equal intervals of wakefulness and sleep (see details above).

### 2.8 Statistical analyses of wake/sleep variables

Statistica 6.1 (StatSoft Inc., Tulsa, OK, USA; RRID: SCR_014213) was used to perform statistical analyses and Prism 10.3 (GraphPad Software Inc., La Jolla, CA, USA; RRID: SCR_002798) to produce figures. Sleep latencies, slopes of recovery, and PS peak frequency were compared between genotypes (WT vs KO) and sex (males vs females) using two-way analyses of variance (ANOVA; with factors Genotype and Sex). Differences in time spent in vigilance state per light/dark periods, per hour, in episode duration, in number of episodes, in time courses of delta, theta, alpha, SW metrics, and multifractal parameters were investigated for BL and REC using three-way repeated-measure ANOVAs with factors Genotype and Sex, and Period, Hour or Interval. Differences in spectral power per Hz-bin were assessed using two-way ANOVAs with factors Genotype and Sex. Differences in SW metrics and multifractal parameters per condition (i.e., BL vs REC) were assessed using three-way ANOVAs with factors Genotype, Sex and Condition. For Figure 1B, ANOVAs were performed including 24 hours for BL and 18 hours for REC, given the lack of variance for hours during the SD. Significant effects in repeated-measure ANOVAs were adjusted using the Huynh-Feldt correction, and significant interactions were decomposed using planned comparisons or Student’s t-tests. The threshold for statistical significance was set to 0.05, and results are reported as mean ± SEM. For analyses of individual electrodes, one WT female and two WT males were removed for the motor cortex and one WT male for the visual cortex because of an artifactual signal for most of the recording of BL, REC or both. One KO female was removed from episode duration for wake under BL condition because it was a significant outlier (Figure 2A; GraphPad ROUT method for outlier detection). One WT female was removed from the 24-hour mean analysis of *Dispersion* and the time course of *Dispersion* during wakefulness, and one WT male was removed from the 24-hour *Dispersion* and the time course of *Dispersion* because of outlier values found for two or more intervals during both days (Figure 5C, 5D and S2D, S2E, S2F;).

**Figure 1.**
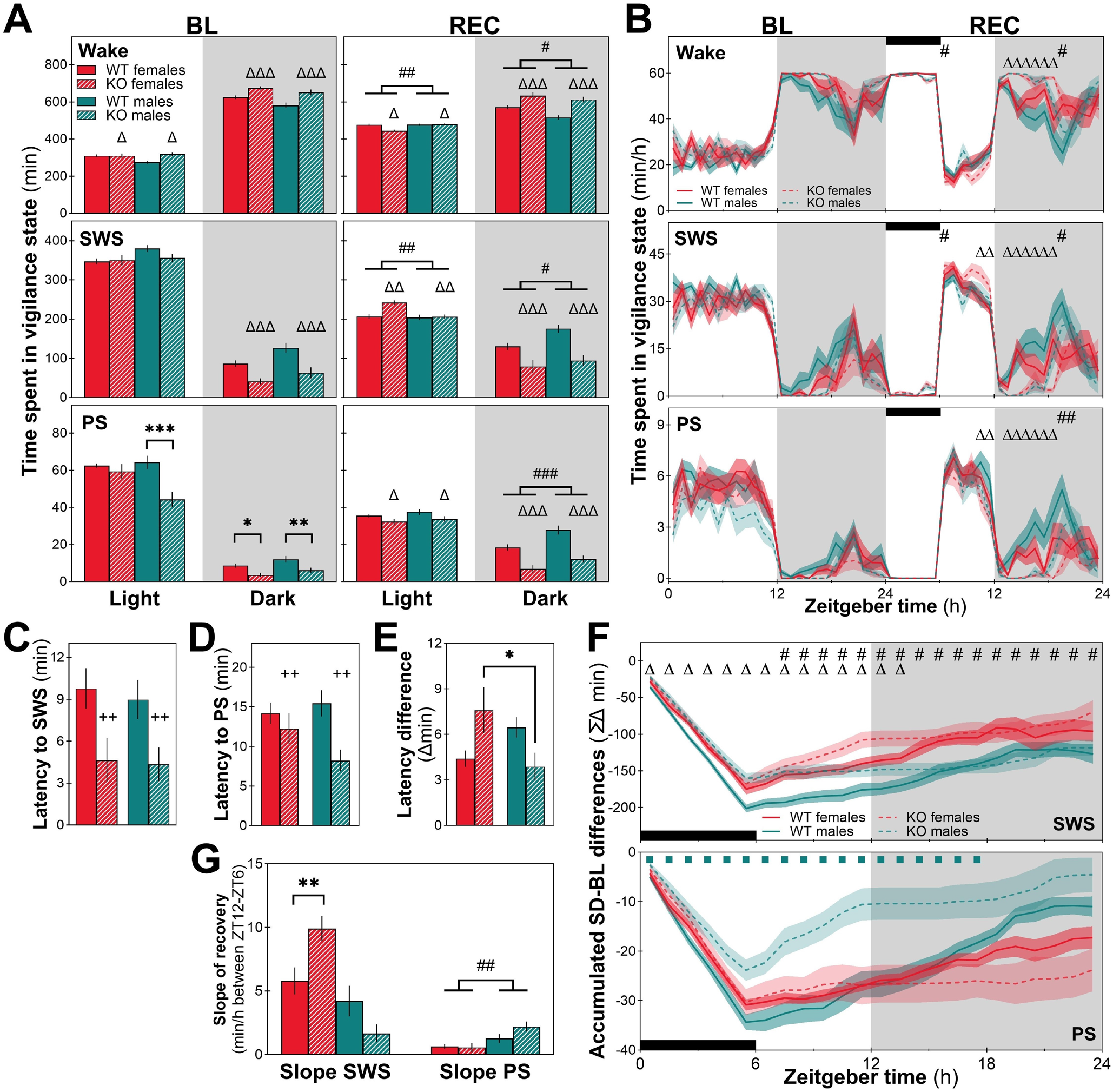
Time spent in wakefulness and sleep states in female and male *Nlgn2* KO mice and WT littermates. **(A)** Time spent in wakefulness, slow wave sleep (SWS) and paradoxical sleep (PS) during the 12-hour light and dark periods of baseline (BL, left column) and sleep deprivation (SD) + recovery (REC, right column). **(B)** Time spent in state per hour in female and male mice of both genotypes. **(C)** Latency to SWS after SD. **(D)** Latency to PS after SD. **(E)** Latency difference between SWS and PS. **(F)** Time course of accumulated differences from BL in SWS and PS calculated from the beginning of the 6-hour SD to 24 hours later. **(G)** Slope of SWS and PS recovery following SD calculated between the 12^th^ and the 6^th^ hour of the 12-hour light period. Differences between WT and KO animals during the 12-hour light/dark periods or at specific hours are represented by Δ: p < 0.05, ΔΔ: p < 0.01, and ΔΔΔ: p < 0.001; differences between males and females during the 12-hour light/dark periods or at specific hours are represented by #: p < 0.05, ##: p < 0.01, and ###: p < 0.001; differences between WT and KO males or WT and KO females or KO females and males are represented by *: p < 0.05, **: p < 0.01, and ***: p < 0.001; differences between WT and KO males at specific hours are represented by turquoise squares: p < 0.05; differences between KO and WT (females and males combined) are represented by ++: p < 0.01. WT females n = 11, WT males n = 9, KO females n = 10, KO males n = 10. Gray backgrounds indicate the 12-hour dark periods. Black rectangles in panels B and C represent the 6-hour SD.

**Figure 2.**
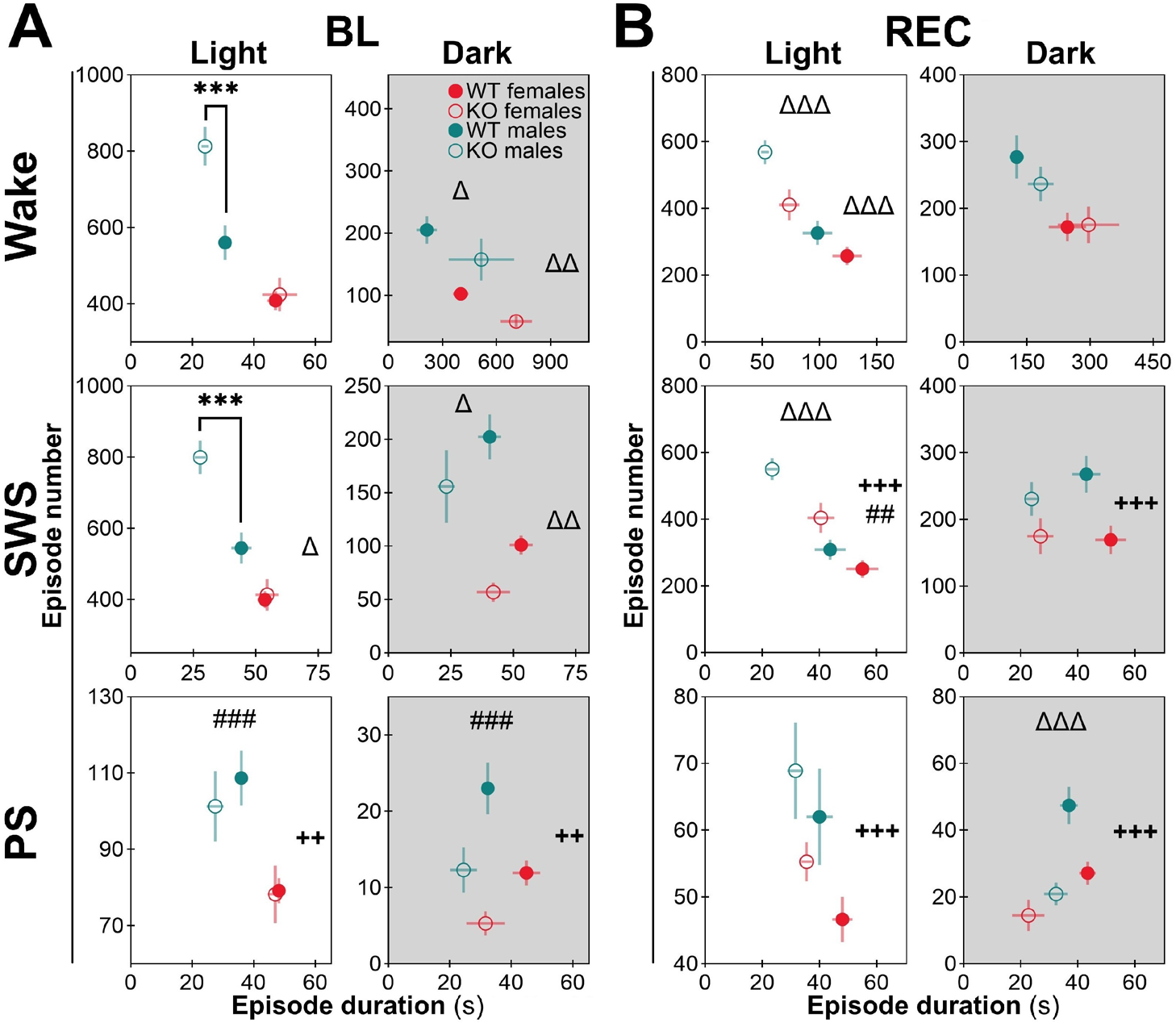
Wake/sleep fragmentation/consolidation in *Nlgn2* KO and WT mice of both sexes for baseline (BL) and recovery (REC). **(A)** Mean duration of individual episodes of wakefulness (x-axis), slow wave sleep (SWS) and paradoxical sleep (PS) in seconds plotted with the number of episodes of each state (y-axis) separately for the BL light and dark periods. **(B)** Same for REC. Symbols on the right relate to episode duration and symbols on the top to episode number; differences between genotypes independent of light/dark periods are represented by ++: p < 0.01, and +++: p < 0.001; differences between genotypes for 12-hour light/dark periods are represented by Δ: p < 0.05, ΔΔ: p < 0.01, and ΔΔΔ: p < 0.001; differences between sexes for the 12-hour light/dark periods are represented by ##: p < 0.01, and ###: p < 0.001; differences between genotypes in males for the 12-hour light period are represented by ***: p < 0.001. WT females n = 11, WT males n = 9, KO females n = 10 (except n = 9 for BL wake in panel A), KO males n = 10. Gray backgrounds indicate the 12-hour dark periods. Upper left quadrant indicative of fragmentation and bottom right quadrant of consolidation.

### 2.9 Transcriptome quantification and analyses

Motor cortex samples of *Nlgn2* KO and WT female and male littermates (WT females n = 4, KO females n = 3, WT males n = 4, KO males n = 3) were submitted to bulk (full tissue) RNA extraction using a PureLink RNA Mini Kit (Thermo Fisher Scientific, Montreal, QC, Canada; #12183018A). The quality of RNA samples was verified using an Agilent 4150 TapeStation System (Agilent Technologies, Santa Clara, CA, USA; G2992AA with RNA screen tape assay; RRID: SCR_019393). For all samples, the RNA integrity number equivalent (RINe) was ≥ 8.2 (concentrations between 149 and 484 ng/μL). Library preparation, sequencing and main analyses were conducted by Genome Québec (Montreal, QC, Canada). Briefly, sequencing was done after preparation of polyA-enriched RNA libraries using a NovaSeq X instrument according to manufacturer’s instructions (PE100; Illumina, San Diego, CA, USA; RRID: SCR_024569). The sequencing resulted in 176 to 270M reads per sample. The DRAGEN RNA v4.3.16 pipeline was used for the alignment of raw FASTQ reads to the *Mus Musculus* (mm10) genome and to exclude low-quality and adapter sub-sequences (alignment rate ≥ 99.3% and unique detected genes ≥ 18,300).

Differentially expressed genes (DEGs) between KO and WT were identified from RNA sequencing data using DESeq2 v1.49.4 (RRID: SCR_015687) separately for females and males. Between-genotype comparisons were computed by fitting a generalized linear model (GLM) with a logarithmic link to the normalized counts while accounting for the gene-wise dispersion. Significance levels obtained from the GLM were adjusted for multiple testing using the Benjamini-Hochberg’s false-discovery rate (FDR). Volcano plots and violin plots were produced with GraphPad Prism 11, and heatmaps using R codes (R Project for Statistical Computing, RRID: SCR_001905; including clustering with ward.D2). A R code was also used to extract DEGs between KO and WT mice that were specific to females, specific to males and shared by males and females. Gene ontology (GO) enrichment analyses interrogating Biological process 2025, Molecular function 2025, Synaptic GO (SynGO) 2024, Reactome pathway 2024, Mouse wikipathway 2024 and Kyoto Encyclopedia of Genes and Genomes (KEGG) pathway 2026 databases were performed with these different ensembles of DEGs (FDR < 0.05) using Enrichr (https://maayanlab.cloud/Enrichr/; Xie et al., 2021). GO terms were considered enriched with enrichment FDR < 0.1 and if including a minimum of two DEGs excluding *Nlgn2*. For the graphical representation, a selection of relevant terms was made and redundant pathways were not depicted. Statistica 6.1 was used to conduct additional two-way ANOVAs with factors Genotype and Sex for specific targets to directly compare the effect of genotype between sexes.

### 2.10 Proteome quantification and analyses

Total proteins (bulk: full tissue) were extracted from the motor cortex of *Nlgn2* KO and WT littermates similar to previously conducted (Areal et al., 2025). Briefly, dissected motor cortices (WT females n = 4, KO females n = 4, WT males n = 5, KO males n = 3) were placed in ice-cold modified RIPA buffer (50 mM HEPES, 20 mM EDTA, 0.1% SDS, 10% IGEPAL, 0.5% sodium deoxycholate, 0.03 mM PMSF, protease inhibitor cocktail [Sigma-Aldrich, St. Louis, MO, USA; #P8340]) and homogenized on ice (Sigma-Aldrich; cordless motor pellet pestle #Z359971). Homogenates were centrifuged at 17,000 g for 10 minutes at 4 °C, and supernatants were kept as the protein extracts. Protein quantification was done using a DC protein assay (Bio-Rad, Montreal, QC, Canada; Reagent A #5000113, Reagent B #5000114, and Reagent S #5000115) with a standard curve of bovine serum albumin (reads done in triplicate at 750 nm), and all protein samples were diluted to 10 μg/μL with ice-cold modified RIPA buffer for proteomic measurement. The quantification of the proteome was conducted by the Proteomic Platform of the Research Institute of the McGill University Health Center using a Thermo Scientific Orbitrap Astral Mass Spectrometer (Thermo Fisher Scientific, Burlington, ON, Canada; RRID: SCR_026205) and a Thermo Scientific Vanquish Neo UHPLC System (RRID: SCR_026495).

A similar number of proteins were detected per sample with 5,304 common between all samples on which the analysis was focused (coefficient of variations within sexes less than 10%). DIA-analyst (https://analyst-suites.org/apps/dia-analyst/; Monash Proteomics and Metabolomics Platform & Monash Bioinformatics Platform, Monash University, Melbourne, Australia) was used for normalization, to calculate peptide imputed intensities, and to extract differentially expressed proteins (DEPs) between KO and WT mice separately for female and males mice (adjusted [adj.] p < 0.05 [t-statistics correction] or p < 0.05 to facilitate enrichment analyses). Volcano plots and heatmaps were produced using R codes (and ward.D2), and box plots with GraphPad Prism. R was also used to extract DEPs that were specific to females, specific to males and shared by males and females using hits that could be statistically compared between genotypes in both sexes. GO enrichment analyses were done as detailed above for the transcriptome using p < 0.05 DEPs (with 2026 Biological process and Molecular function databases since updated after transcriptomic analyses; selection of relevant terms for figures and redundant pathways not depicted). Statistica 6.1 was used to conduct two-way ANOVAs (factors Genotype and Sex) for selected targets to directly compare the effect of genotype between sexes. Lastly, an integration of the transcriptome and proteome was conducted by calculating the number of hits that overlap between DEGs (FDR < 0.05) and DEPs (p < 0.05). This was done by merging DEGs and DEPs found in female and male mice.

## 3 Results

### 3.1 Less time spent asleep in *Nlgn2* KO females and males

The time spent in vigilance states was initially compared between *Nlgn2* KO mice and WT and between sexes over an undisturbed 24-hour day (BL). *Nlgn2* KO animals spent significantly more time awake than WT animals, especially during the 12-hour dark period, and less time in SWS only during the 12-hour dark period (Figure 1A, upper and middle left panels; Table S1A). Also, *Nlgn2* KO males spent less time in PS than WT males during the 12-hour light and dark periods of BL, while the difference was only significant for the dark period in *Nlgn2* KO females (Figure 1A, lower left panel; Table S1A). The analysis of the 24-hour distribution of time spent in states revealed that the abovementioned differences are not linked to specific hours of the BL light and dark periods but are rather global (coming from the combination of several hours; Figure 1B; Table S1B).

The response to a homeostatic sleep challenge was then interrogated by comparing genotypes and sexes for time spent in states during a 6-hour SD and the following 18 hours of undisturbed recovery. Under these conditions (REC), *Nlgn2* KO animals of both sexes spent less time awake and more time in SWS during the light period, but more time awake and less in SWS during the dark period when compared to WT littermates (Figure 1A, upper and middle right panels; Table S1A). Similar to males during BL, animals lacking NLGN2 spent less time in PS during the light and dark periods of REC (Figure 1A, lower right panel; Table S1A). These genotype differences were mainly driven by an increased time spent in SWS in KO compared to WT and a decreased time spent in PS at the end of the REC light period, and by more time spent awake and less time spent in SWS and PS during the first half of the REC dark period (Figure 1B; Table S1B).

In addition, main effects of sex on the time spent in states were observed specifically when considering REC (i.e., SD and recovery). Indeed, females spent less time awake and more time in SWS in comparison to males during the light period (significant for the first hour of recovery after SD), but more time awake and less time in SWS during the dark period of REC (mainly middle of the dark period; Figure 1A right panels and Figure 1B; Table S1A and S1B). Moreover, females spent less time in PS than males during the dark period of REC (mainly middle of the dark period; Figure 1A, lower right panel and Figure 1B; Table S1A and S1B). A significant effect of sex on time spent in SWS after elevated homeostatic sleep pressure was also reported in a previous study (Paul et al., 2006).

Following SD, females and males lacking NLGN2 fell asleep significantly faster than WT animals with latencies to reach SWS and PS being significantly shorter (Figure 1C and Figure 1D; Tables S1C and S1D). Interestingly, KO females expressed a delay in reaching PS after SWS initiation in comparison to KO males (Figure 1E; Table S1E). Accumulated differences were computed to interrogate SWS and PS loss during SD and their recovery following SD (when compared to BL). For SWS, *Nlgn2* KO females and males accumulated less difference from BL in comparison to WT mice from the beginning of SD to the second hour of the dark period (Figure 1F, upper panel; Table S1F). Also, females recovered more SWS than males from the second hour after the end of SD to the end of the dark period (Figure 1F, upper panel; Table S1F). For PS, only *Nlgn2* KO males accumulated less difference from BL than WT males between the beginning of SD until the middle of the dark period (Figure 1F, lower panel; Table S1F; female KO mice not significantly different from female WT mice). These results suggest differences in the recovery process of KO animals, which was further interrogated by comparing the slope of initial recovery (between ZT6 and ZT12) for SWS and PS. During the light period of REC, females lacking NLGN2 recovered SWS almost two times faster than WT females, while there was no significant difference between KO and WT in males (Figure 1G; Table S1G). For PS, there was no significant difference between KO and WT females/males, but males recovered significantly faster than females during the first 6 hours of recovery (Figure 1G; Table S1G). Overall, these results suggest that *Nlgn2* KO females recover SWS faster due to enhanced SWS after SD, while KO males recover PS faster, which could result from less time spent in PS during BL (e.g., greater opportunity to increase time in PS immediately after SD given a ‘naturally’ reduced time in this state at this time of day).

### 3.2 Sleep fragmentation particularly in *Nlgn2* KO males

Wake/sleep fragmentation/consolidation was investigated using the mean duration of individual episodes and the number of episodes of the three vigilance states (in Figure 2, fragmentation indicated by values closer to the upper left quadrant and consolidation by values approaching the bottom right quadrant). Regarding BL mean episode duration, *Nlgn2* KO females and males had longer episodes of wakefulness specifically during the dark period when compared to WT littermates (Figure 2A, x-axis of upper panels; Table S2A; no difference during the light period). Moreover, *Nlgn2* KO displayed shorter SWS episode duration (significant for both the light and dark periods; Figure 2A, x-axis of middle panels; Table S2A), and shorter PS episode duration (significant considering light and dark periods altogether; Figure 2A, x-axis of lower panels; Table S2A). For the number of episodes of vigilance states under BL, only *Nlgn2* KO males showed more wake and SWS episodes during the light period in comparison to WT males (no difference between KO and WT in females; Figure 2A, y-axis of upper and middle panels; Table S2B). During the BL dark period, *Nlgn2* KO females and males showed less wake and SWS episodes in comparison to WT littermates (Figure 2A, y-axis of upper and middle panels; Table S2B). In parallel, independently of genotype, male mice expressed more PS episodes than females during both the light and dark periods of BL (Figure 2A, y-axis of bottom panels; Table S2B).

During REC, a reduction of the mean duration of episodes was observed in mice lacking NLGN2 in comparison to WT mice, and this difference was specific to the light period for wake and encompassed both the light and dark periods for SWS and PS (Figure 2B, x-axis of left and right panels; Table S2A). In parallel, female mice had a significantly longer duration of SWS episodes than males during the light period of REC independently of genotype (Figure 2B, x-axis of middle left panel; Table S2A). Under REC conditions, *Nlgn2* KO females and males also displayed a higher number of wake and SWS episodes during the light period, and a lower number of PS episodes during the dark period in comparison to WT mice (Figure 2B, y-axis; Table S2B). In general, these results suggest a fragmentation of wake and SWS states in the absence of NLGN2 when considering the main rest period (i.e., light) that is manifest under both BL and challenged condition in males, but only under challenged conditions in females, as well as a consolidation of wakefulness during the BL active (dark) period.

### 3.3 Altered ECoG spectral power in *Nlgn2* KO females and males

To determine the influence of NLGN2 on the quality of wake/sleep states in male and female mice, the ECoG power spectrum (0.75 to 50 Hz) was compared between genotypes and sexes for each vigilance state, separately for BL and REC. For wakefulness, *Nlgn2* KO mice displayed more spectral power for most Hz-bins from 0.75 to 34 Hz during BL and REC, and also less activity above 46.5 Hz during BL. Under BL, females (independently of genotype) presented higher power in wake for most frequency bins between 8 and 10 Hz and between 28 and 45 Hz, encompassing high theta and low gamma ranges (Figure 3A, upper panels; Table S3A-1). For SWS, animals lacking NLGN2 showed more power over the entire spectrum under both BL and REC (Figure 3A, middle panels; Table S3A-2). For PS, *Nlgn2* KO males had significantly higher power than WT males for most Hz-bins between 3.75 to 6.25 Hz and between 11.75 to 25 Hz during BL, encompassing high delta/low theta and beta ranges, which was also found for frequencies between 4 and 5.25 Hz and between 10 and 25.75 Hz during REC (Figure 3A, lower panels; Table S3A-3). These differences in PS spectra were significantly larger in *Nlgn2* KO females that showed higher power than WT females for most frequencies between 3.75 to 6.25 Hz and between 11.75 to 26 Hz during BL, and between 4 and 5.25 Hz and between 10 and 27.5 Hz during REC (Figure 3A, lower panels; Table S3A-3). The theta peak frequency of PS was also computed to further quantitatively describe the spectral signature of *Nlgn2* KO mice. During both BL and REC, the lack of NLGN2 induced a shift of PS peak frequency towards slower frequencies that was similar in females and males (KO mice having a peak frequency around 6 Hz and WT animals around 7.25 Hz; Figure 3A, insets of lower panels; Table S3A-4). Overall, the absence of NLGN2 in mice induced substantial alterations in spectral ECoG activity in all three vigilance states, with indications that females are more affected by the absence of this synaptic protein when considering PS. Of importance is that the widespread effects of the mutation on vigilance state power spectra are also observed when considering the motor and visual cortex separately (Supplementary figure 1).

**Figure 3.**
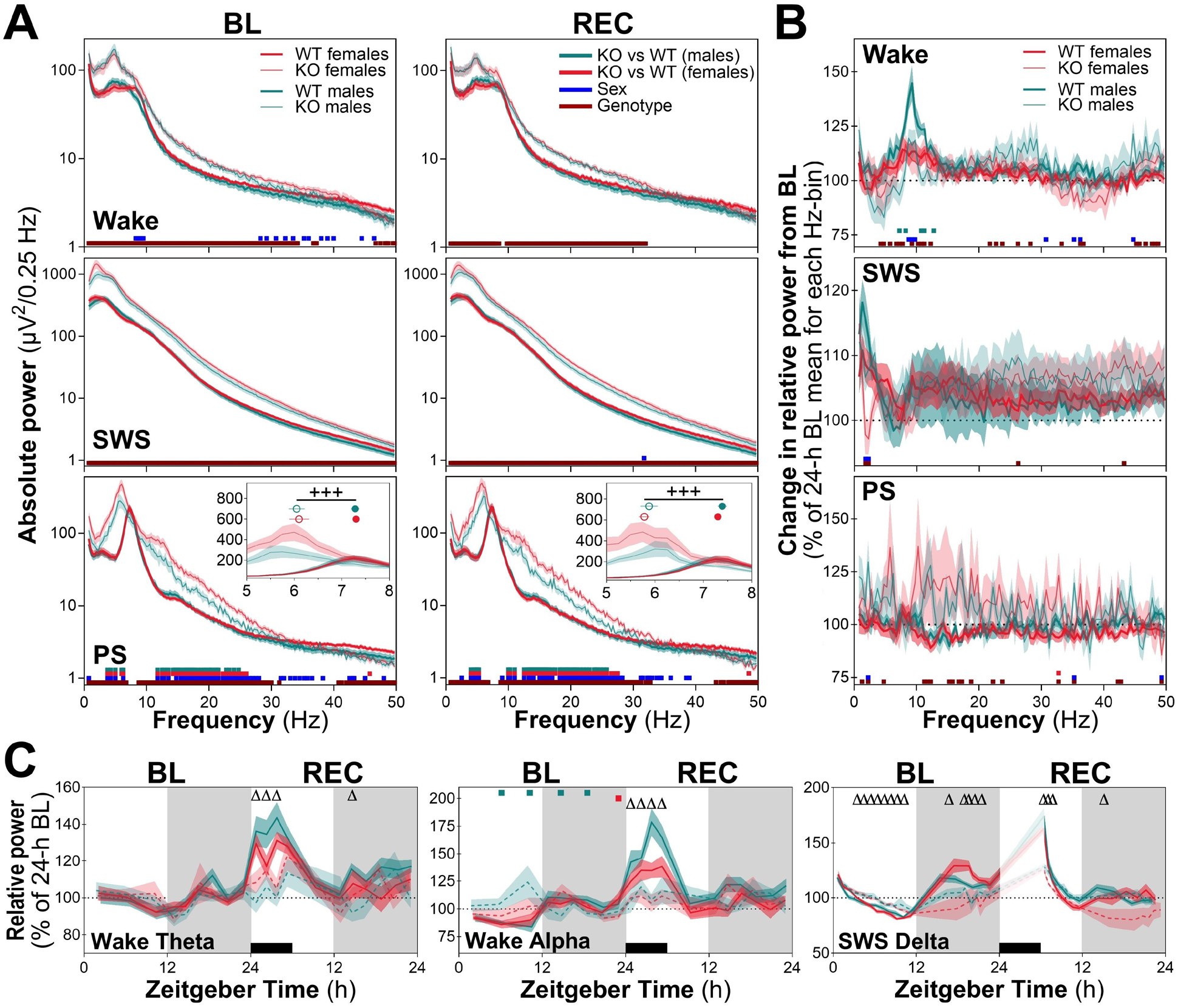
ECoG spectral power for wakefulness, slow wave sleep (SWS), and paradoxical sleep (PS) under baseline (BL) and recovery (REC) and spectral activity time courses for three frequency bands in *Nlgn2* KO and WT males and females. **(A)** Absolute spectral power (log scale) between 0.75 and 50 Hz for wakefulness, SWS, and PS for the 24-hour BL and REC. Insets in PS panels show the theta peak frequency. **(B)** Relative spectral power during REC for frequencies between 0.75 and 50 Hz for wake, SWS, and PS expressed as a percentage of BL. **(C)** 24-hour time courses of relative wake theta (6-9 Hz; left panel), wake alpha (9-12 Hz; middle panel), and SWS delta (1-4 Hz; right panel) activity during BL and REC. Genotype differences per 0.25 Hz-bin are represented by burgundy bars above the x-axes (p < 0.05); sex differences are represented by dark blue bars (p < 0.05); differences between WT and KO males or WT and KO females are represented by turquoise or red bars, respectively (p < 0.05); genotype differences at specific intervals are represented by Δ: p < 0.05; genotype main effects are represented by +++: p < 0.001. WT females n = 11, WT males n = 9, KO females n = 10, KO males n = 10. PS peak frequency is displayed as open (KO mice) and closed (WT mice) circles and red and turquoise represent, respectively, female and male animals. In panel C, gray backgrounds indicate the 12-hour dark periods and black rectangles represent the 6-hour SD.

Next, the effects of the mutation on the ECoG power response to SD (i.e., changes from BL) was verified by comparing genotypes and sexes for the 24-hour relative power of REC expressed as percent change from the 24-hour BL power. For wake, *Nlgn2* KO males were lacking the increase in spectral power for most Hz-bins between 7.25 and 12.75 Hz that was observed for WT males during REC (Figure 3B, upper panel; Table S3B). Moreover, *Nlgn2* KO females and males showed changes from BL that were not found in WT littermates, such as an increased wake power for some frequencies between 21.75 and 28.25 Hz and above 45 Hz (encompassing beta and low gamma ranges), and a reduced wake power for some bins between 33.25 and 36.75 Hz (Figure 3B, upper panel; Table S3B). For SWS, significant genotype and sex differences were found for spectral power change from BL mainly driven by the observation of *Nlgn2* KO females having a decreased ECoG power change between 1.75 and 2.25 Hz (Figure 3B, middle panel; Table S3B). For PS, *Nlgn2* KO females and males showed an enhanced power relative to BL for many Hz-bins encompassing the full interrogated spectrum (Figure 3B, lower panel; Table S3B). Globally, the effects of the absence of NLGN2 on the ECoG power change between REC and BL was dependent on sex, in particular concerning wakefulness.

To unveil the 24-hour dynamics of spectral activity in specific frequency bands, the time courses of wake theta (6-9 Hz), wake alpha (9-12 Hz), and SWS delta (1-4 Hz) relative activity were analyzed separately for BL and REC. Regarding theta activity during wakefulness, no significant difference was found between genotypes and sexes during BL. However, *Nlgn2* KO females and males (combined sexes) showed a major attenuation of the increase in theta activity occurring during SD that was observed in WT animals, which also applied to wake alpha activity (Figure 3C, left and middle panels; Table S3C). Concerning the 24-hour BL dynamics of wake alpha activity, *Nlgn2* KO males showed higher activity than WT males for some intervals during the light period, but a decreased activity for intervals during first half of the dark period (Figure 3C, middle panel; Table S3C). Finally, during BL, *Nlgn2* KO females and males (combined sexes) showed a generally lower amplitude of the 24-hour variations in SWS delta activity when compared to WT littermates, with relative delta activity being higher for most intervals of the light period, but lower for most intervals of the dark period (Figure 3C, right panel; Table S3C). During REC, *Nlgn2* KO females and males showed lower delta activity than WT mice immediately after the end of SD and at the beginning of the dark period (Figure 3C, right panel; Table S3C). This last observation suggests a reduced rebound of SWS intensity after SD in the absence of NLGN2, which is independent of sex.

### 3.4 Higher slow wave amplitude and slope in *Nlgn2* KO females and males

The density and some properties of individual SW during SWS, including amplitude, slope, positive phase duration and negative phase duration, were compared between genotypes and sexes as these represent additional homeostatically-regulated proxies of sleep intensity (Freyburger et al., 2017; Massart et al., 2014; Mongrain et al., 2011). The 24-hour mean number of SW per minute of SWS, or density, was higher in *Nlgn2* KO mice in comparison to WT littermates under both BL and REC, independent of sex (Figure 4A, upper panel; Table S4A). This difference was particularly prominent during the BL light period, and the light and dark periods of REC (Figure 4B, upper panel; Table S4B). The amplitude and slope of SW were both higher in mice lacking NLGN2 in comparison to WT littermates during BL and REC, differences that were not affected by sex and not specific to any intervals of the 24-hour days (Figure 4A and Figure 4B, second and third rows; Table S4A and S4B). Negative phase duration was shorter in *Nlgn2* KO than in WT mice for both BL and REC with again no influence of sex, and with significant differences found for all intervals of BL and most intervals of REC (Figure 4A and Figure 4B, fourth row; Table S4A and S4B). Finally, the 24-hour mean duration of the positive phase of SW and its 24-hour dynamics during BL and REC were not significantly affected by the absence of NLGN2 (Figure 4A and Figure 4B, fifth row; Table S4A and S4B). Overall, these results highlight that the lack of NLGN2 in mice induces multiple alterations of SW density/properties during SWS, and that these differences are not shaped by the biological sex of the animals. Biological sex in itself also showed only a minimal impact on SW density and properties quantified during BL and REC (Figure 4 and Tables S4).

**Figure 4.**
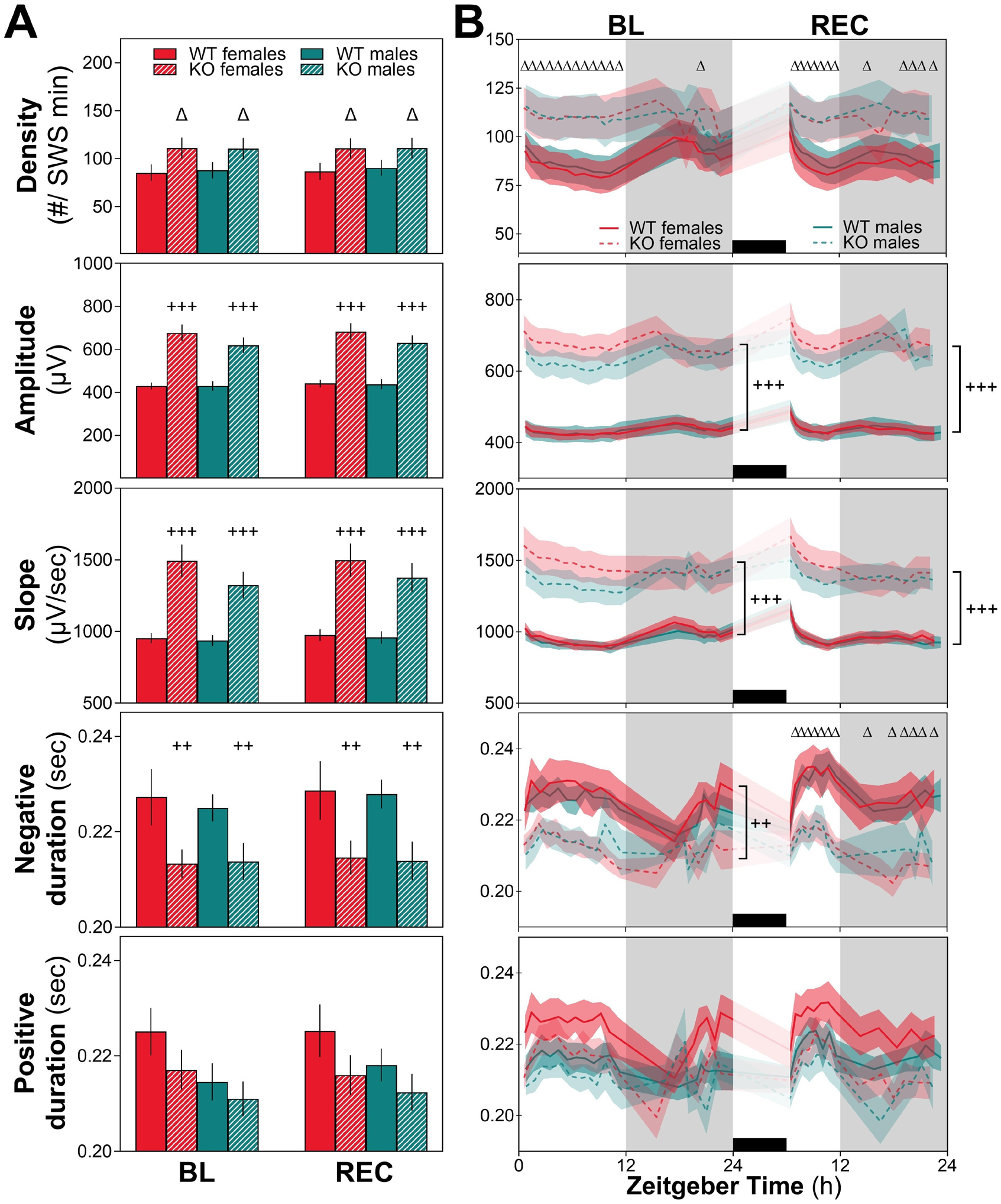
Density and properties of slow waves (SW) under baseline (BL) and recovery (REC) in *Nlgn2* KO and WT females and males. **(A)** SW density and properties (amplitude, slope, negative phase duration, positive phase duration) averaged for the 24-hour BL and the 24-hour REC. **(B)** 24-hour time courses during BL and REC of SW density and properties averaged across intervals comprising the same number of SWS epochs. Differences between WT and KO mice encompassing BL and REC in A panels or all intervals of BL or REC in B panels are represented by ++: p < 0.01, and +++: p < 0.001; differences between WT and KO animals found separately for BL and REC (A panel) or at specific intervals (B panels) are represented by Δ: p < 0.05. WT females n = 11, WT males n = 9, KO females n = 10, KO males n = 10. Gray backgrounds indicate the 12-hour dark periods, and black rectangles represent the 6-hour sleep deprivation (SD).

### 3.5 Altered aperiodicity metrics in *Nlgn2* KO females and males

Spectral analysis provides information about brain rhythmic activity but lacks information regarding the arrhythmic/aperiodic component of brain activity. A multifractal analysis was used to assess arrhythmic ECoG properties and the long-range relationships in the activity across different time scales (Hu et al., 2013; Lina et al., 2019). Independent of sex, *Nlgn2* KO mice were found to have higher *Hm* values for wake, SWS and PS for both BL and REC (Figure 5A and Figure 5B, Supplementary figure 2A; Tables S5A and SF2). For wakefulness, this difference appeared to predominate during light periods (Figure 5B, upper panel; Table S5B). For SWS, the higher *Hm* was global over both 24-hour days in KO animals when compared to WT littermates (Figure 5B, second panel; Table S5B). For PS, the impact of the mutation on *Hm* during BL was influenced by time-of-day and by sex, with *Nlgn2* KO females showing larger differences from WT, especially at the beginning of the dark period (Figure 5B, lower panel; Table S5B). During REC, PS *Hm* was higher in the absence of NLGN2, particularly at the end of the light period and independently of sex. Overall, these data suggest that *Nlgn2* KO females and males equally display more persistence in the long-range temporal dependency of ECoG activity in all three vigilance states. It is interesting to note that these differences are also manifest when examining the motor and visual cortex separately (Supplementary figure 2 and Table SF2).

**Figure 5.**
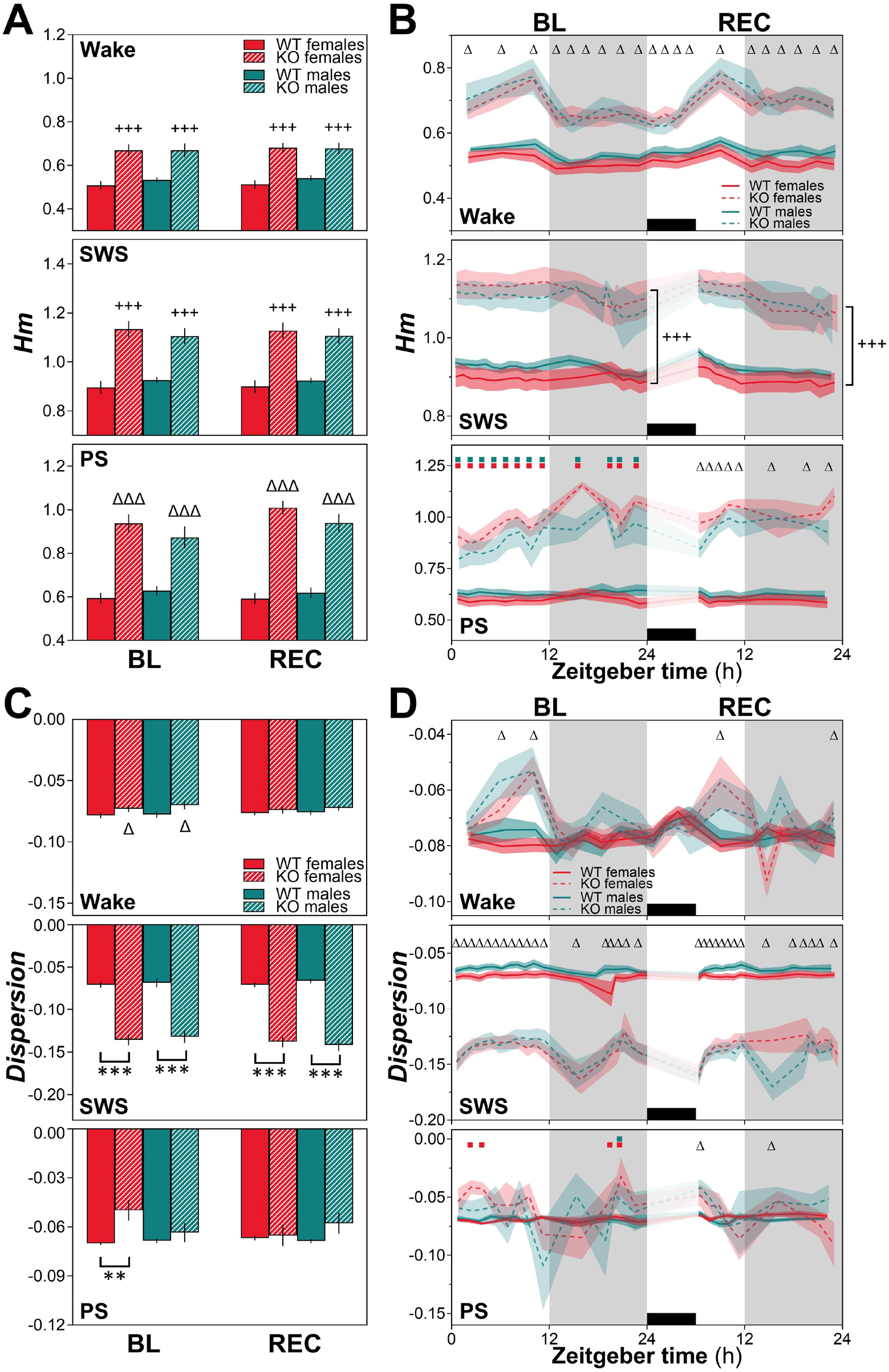
The most prevalent Hurst exponent (*Hm*) and *Dispersion* of exponents around *Hm* computed for wakefulness, SWS, and PS during baseline (BL) and recovery (REC) in *Nlgn2* KO and WT females and males. **(A)** *Hm* values averaged over 24 hours for BL and REC. **(B)** 24-hour time courses of *Hm* for the three vigilance states. **(C)** *D* values averaged over 24 hours for BL and REC. **(D)** 24-hour time courses of *Dispersion* for the three vigilance states. Differences between WT and KO mice encompassing BL and REC and all intervals of BL or REC are represented by +++: p < 0.001; differences between WT and KO mice that differ between BL and REC or between intervals are represented by Δ: p < 0.05 and ΔΔΔ: p < 0.001; differences between WT and KO males or WT and KO females that are impacted by BL/REC are represented by **: p < 0.01 and ***: p < 0.001; differences between WT and KO males or WT and KO females at specific intervals are represented by turquoise and red squares, respectively (p < 0.05). WT females n = 11 (except n = 10 for wake in panels C and D), WT males n = 9 (except n = 8 for panels C [wake and PS] and D), KO females n = 10, KO males n = 10. Gray backgrounds indicate the 12-hour dark periods, and black rectangles the 6-hour sleep deprivation (SD).

For *Dispersion* during wakefulness, *Nlgn2* KO females and males (combined sexes) had less negative values for BL (Figure 5C, upper panel; Table S5C), which was driven by genotype differences restricted to the light period (Figure 5D, upper panel; Table S5D). For REC, less negative values of wake ECoG *Dispersion* were found only for two intervals across the recovery after SD (Figure 5D, upper panel; Table S5D). Conversely, the *Dispersion* of the SWS ECoG was more negative in mice lacking NLGN2 under both BL and REC (Figure 5C, second panel; Table S5C), with the magnitude of the genotype difference being significantly affected by time-of-day but not by sex (Figure 5D, second panel; Table S5D). For *Dispersion* of the PS BL signal, only *Nlgn2* KO females showed a less negative value in comparison to WT females (Figure 5C, bottom panel; Table S5C) although KO males showed a similar difference when considering separately the BL light and dark periods (Supplementary figure 2D and Table SF2). The genotype difference in females appeared mainly driven by significant differences during the first half of the light period and the last half of the dark period (Figure 5D, bottom panel; Table S5D). During REC, the difference between KO and WT was in the same direction (less negative), and found to be significant for only one interval per light/dark period (Figure 5D, bottom panel; Table S5D). These results suggest that *Nlgn2* KO mice display differences from WT mice in the multifractal profile of the ECoG that are dependent on the vigilance state, with indication for less multifractality during wake and PS (particularly evident for females in PS), but for more multifractality during SWS. These differences are likely driven by alterations in the crosstalk/connectivity between brain regions, especially since KO mice show changes in the opposite direction during PS when considering separately the signal of the motor and visual cortex (Supplementary figure 2E and 2F).

### 3.6 Sex-specific transcriptome of *Nlgn2* KO mice

To get indications on molecular and cellular pathways and functions contributing to the major alterations of ECoG spectral and aperiodic profiles of *Nlgn2* KO females and males, genome-wide gene expression was measured for the motor cortex (bulk measurement: all tissue cells). In females, 112 DEGs were found in KO compared to WT animals (including *Nlgn2*), with a roughly similar proportion showing a lower (45.5%) or a higher (54.5%) expression in KO (Figure 6A, upper panel; Table S6B). In males, 311 DEGs were observed in KO mice with more than 64% showing a higher expression in mutants (Figure 6A, lower panel; Table S6C). From these, 34 were found to be significantly changed in both sexes (all in the same direction in females and males), including a majority (76.5%) that were expressed at a higher level in mutants (Figure 6B; Table S6D). These 34 DEGs in KO females and males were found to be enriched notably for genes related to MAPK/ERK and G protein-coupled receptor (GPCR) signaling pathways, protein regulation (e.g., secretion, phosphorylation, degradation), and neurotransmission (e.g., neuropeptide activity, chemical synaptic transmission; Figure 6C and Table S6E). Selected representative examples of genes are shown in Figure 6D (*Pnoc* for neurotransmission; *Wnt2b* and *Pthlh* for GPCR signaling; *Adamts8* and *Scg2* for protein regulation/metabolism; *Ret* for MAPK signaling). *Npy* and *Cort* also represent hits commonly higher in expression in KO females and males (*Npy* among those with highest significance level) that associate to neurotransmission and GPCR signaling (Figure 6A).

**Figure 6.**
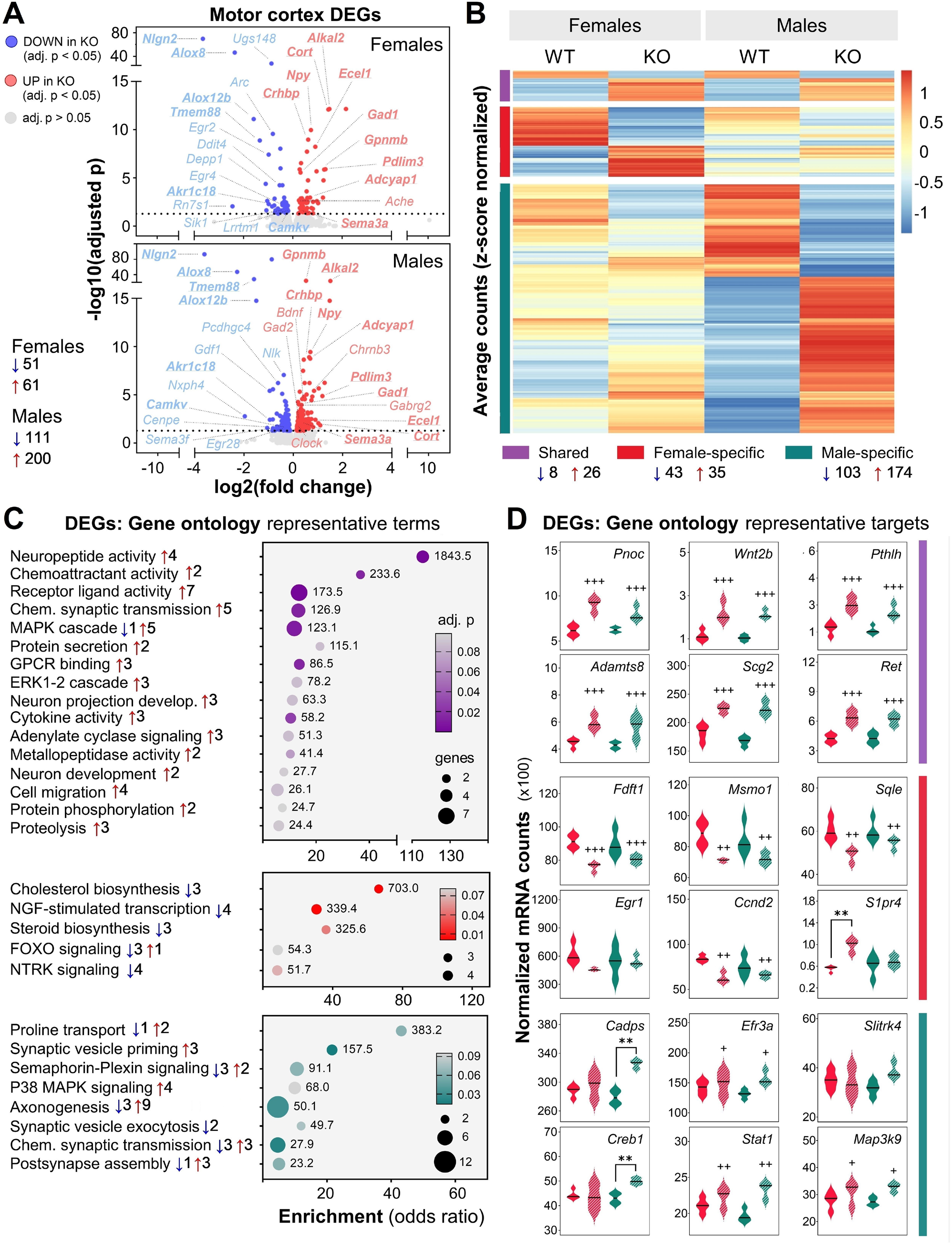
Motor cortex transcriptome of *Nlgn2* KO female and male mice compared to that of WT littermates. **(A)** Volcano plots for genotype comparisons in females (top panel) and males (bottom panel), with numbers of decreased and increased differentially expressed genes (DEGs: FDR < 0.05) in KO mice indicated on the bottom left. Gene symbols are indicated for some of the top DEGs, with underlined and bold symbols indicative of DEGs found in both sexes. **(B)** Heatmap showing DEGs that are shared between sexes (purple), or that are specific to females (red) and males (green). Average normalized counts per group were z-scored by gene to emphasize differences. **(C)** Representative gene ontology terms obtained using the three different DEG clusters of panel B. An adjusted (adj.) p value < 0.1 was used as threshold for the representation of enriched terms. Numbers of increased and decreased DEGs in KO relative to WT mice are specified next to each term. Dot size and color respectively represent the number of genes and the significance. Scores of combined enrichment and significance metrics are written next to each dot. **(D)** Expression of representative DEGs selected amongst the enriched gene ontology terms separately for each cluster. Females are indicated by red/pink violin and males by green ones, with solid colors representing WT mice and hatched ones KO mice. Differences between KO and WT (females and males combined) are represented by +: p < 0.05, ++: p < 0.01 and +++: p < 0.001. Differences between WT and KO males or between WT and KO females are represented by **: p < 0.01. WT females n = 4, KO females n = 3, WT males n = 4, KO males n = 3.

A total of 78 DEGs between KO and WT were found only in females (female-specific DEGs), which include 55.1% with expression being lower and 44.9% being higher in mutants (Figure 6B). These were found to be enriched in particular for genes related to cholesterol/steroid synthesis (e.g., *Fdft1*, *Msmo1*, *Sqle*) as well as NTRK (e.g., *Arc*, *Egr1*) and FOXO (e.g., *Ccnd2*, *S1pr4*) signaling pathways (Figure 6A, Figure 6C and Figure 6D; Table S6F). However, it is important to note that > 80% of female-specific DEGs were not captured by the gene ontology analysis (Table S6B and Table S6F). Moreover, even if the gene expression changes did not reach genome-wide statistical significance in males, a significant difference between *Nlgn2* KO and WT male mice can still be observed for many of these hits (even if generally of lower magnitude) when gene expression levels are directly compared between sexes and genotypes separately for each hit (i.e., *Ccnd2*, *Msmo1*, *Sqle*, *Fdft1*; Figure 6D). When considering genotype differences only reaching genome-wide significance in males, 277 DEGs were found (62.8% higher in KO than WT; 3.5 times more than in females; Figure 6B; Table S6C). This cluster was enriched, among others, in genes related to synaptic vesicle regulation (*Cadps*, *Efr3a*), as well as signaling-related protein phosphorylation and transcriptional regulation (P38 MAPK linked genes: *Creb1*, *Stat1*, *Map3k9*; Figure 6C and Figure 6D; Table S6G). Once more, some of these male-specific hits seem to show some level of genotype difference also in females (Figure 6D). *Slitrk4* was also depicted in Figure 6D (linked to axonogenesis, Table S6C) because of its proposed role in neurodevelopmental disorders (Amelan et al., 2026). Globally, despite some similarities between sexes in affected pathways, NLGN2 absence appears to drive more extensive changes to the motor cortex transcriptome of males in comparison to that of female mice, with a relatively low proportion of common genes when considering the conservative genome-wide correction (10-30%).

### 3.7 Sex-specific proteome of *Nlgn2* KO mice

To further interrogate functional elements with potential to contribute to *Nlgn2* KO female/male ECoG spectral and aperiodic profiles, the proteome was measured also for the motor cortex (bulk measurement: all tissue cells). In females, 33 DEPs were found in KO compared to WT animals when using proteome-wide correction (including NLGN2), with 63.6% showing a lower level (e.g., SHANK1) and 36.4% a higher level (e.g., THY1) in KO (Figure 7A, upper panel; Table S7B). In males, 19 DEPs were observed in KO mice with about half showing a higher level (e.g., PVALB) in mutants (Figure 7A, lower panel; Table S7C). To facilitate the identification of shared and sex-specific changes in protein level and of associated molecular and cellular pathways, DEPs (p < 0.05) found without proteome-wide correction were then considered (i.e., 400 in females and 398 in males, 53.2-55.5% higher in KO; Figure 7A). From these and using only proteins with statistical comparison available for both sexes, 56 were found to be changed by genotype in both sexes (7 in different directions in females and males), including 48.2% that were expressed at a lower level in mutants of both sexes (Figure 7B; Table S7D). These 56 DEPs in KO females and males were found to be enriched in protein-coding genes related notably to calcium signaling (e.g., CACNA1E, SLC8A2), excitatory synapse function (ERK2/MAPK1, *Grm2*/mGluR2, *Grm5*/mGluR5) and neurodegenerative disease and oxidative reaction (COX7A1) (Figure 7C and Figure 7D; Table S7E). PKCγ (*Prkcg*) was also found to be a key protein commonly lower in KO females and males (corrected p < 0.05) that associates to calcium signaling and excitatory synapse function (Figure 7A; Table S7E).

**Figure 7.**
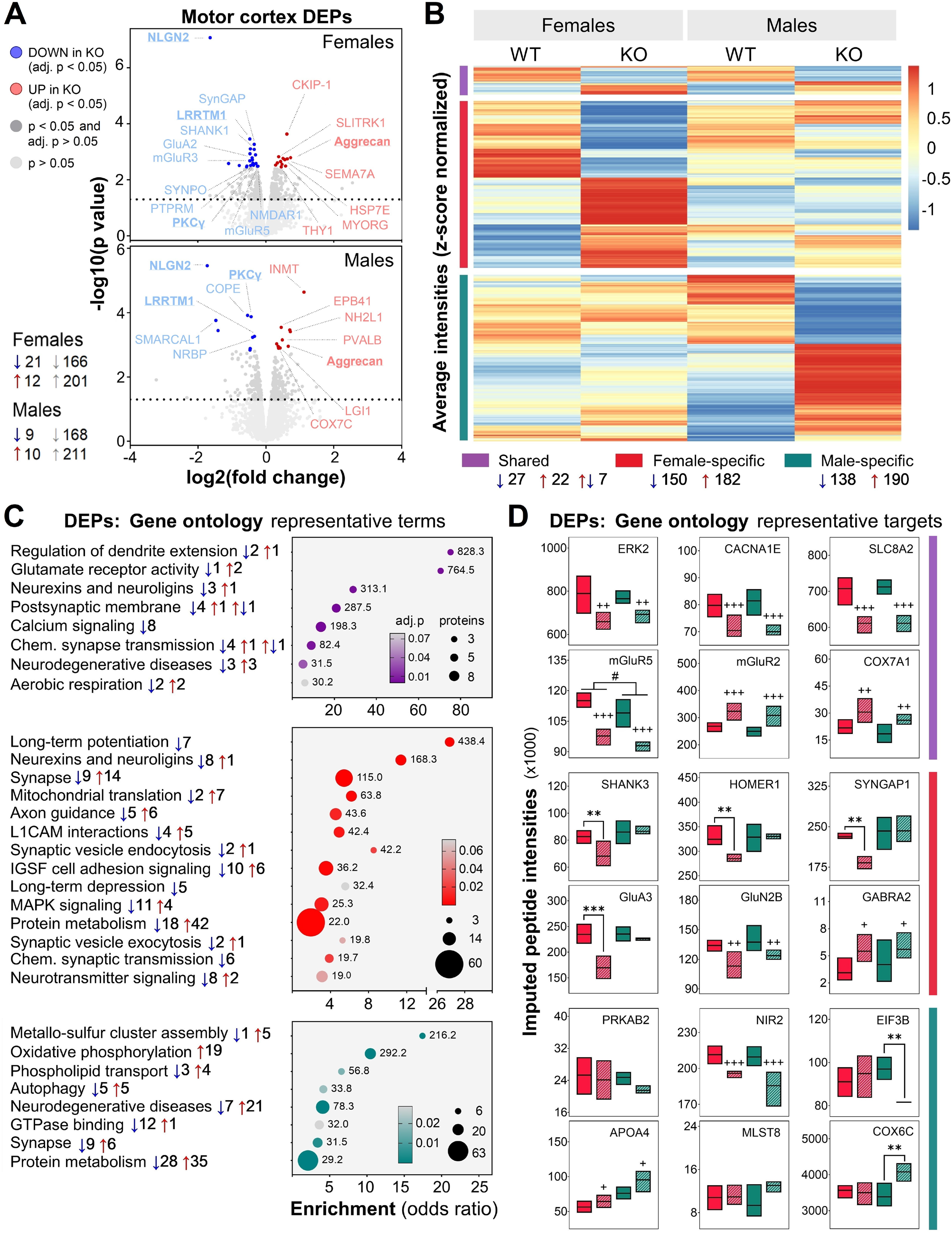
Motor cortex proteome of *Nlgn2* KO female and male mice compared to that of WT littermates. **(A)** Volcano plots for genotype comparisons in females (top panel) and males (bottom panel), with numbers of decreased and increased differentially expressed proteins (DEPs) in KO mice indicated on the bottom left (adjusted [adj.] p < 0.05 shown with blue/red colors and p < 0.05 shown in darker grey). Protein symbols are indicated for some of the top DEPs, with underlined and bold symbols indicative of DEPs found in both sexes. **(B)** Heatmap showing DEPs (p < 0.05) that are shared between sexes (purple), or that are specific to females (red) and males (green). Average imputed intensities per group were z-scored by protein to emphasize differences. **(C)** Representative gene ontology terms obtained using the three different DEP clusters of panel B. An adj. p value < 0.1 was used as threshold for the representation of enriched terms. Numbers of increased and decreased DEPs in KO relative to WT mice are specified next to each term. Dot size and color respectively represent the number of proteins and the significance. Scores of combined enrichment and significance metrics are written next to each dot. **(D)** Expression of representative DEPs selected amongst the enriched gene ontology terms separately for each cluster. Females are indicated by red/pink boxes and males by green ones, with solid colors representing WT mice and hatched ones KO mice. Differences between KO and WT (females and males combined) are represented by +: p < 0.05, ++: p < 0.01 and +++: p < 0.001, and the only difference between females and males is represented by #: p < 0.05. Differences between WT and KO males or between WT and KO females are represented by **: p < 0.01 and ***: p < 0.001. WT females n = 4, KO females n = 4, WT males n = 5, KO males n = 3.

An ensemble of 332 DEPs between KO and WT were found only in females (female-specific DEPs), which include 45.2% with level being lower and 54.8% being higher in mutants (Figure 7B; Table S7D). These were found to be enriched for protein-coding genes related, in particular, to excitatory and inhibitory synaptic function and plasticity (e.g., SHANK3, HOMER1, SYNGAP1, *Gria3*/GluA3, *Grin2b*/GluN2B, *Gabra2*/GABAARα2), and also to mitochondrial functions (e.g., MRPL3, MRPL9) (Figure 7C and Figure 7D; Table S7F). However, once again for specific hits, even if the protein level changes did not reach proteome-wide statistical significance in males, a significant difference between *Nlgn2* KO and WT male mice can still be observed (even if lower in magnitude) when protein levels are directly compared between sexes and genotypes separately for each hit (i.e., GluN2B, *Gabra2*/GABAARα2; Figure 7D). When considering genotype differences only significant in males, 328 DEPs were found (57.9% higher in KO than WT; Figure 7B; Table S7D). This cluster was found to be enriched, among others, for protein-coding genes related to autophagy (e.g., PRKAB2, MLST8), phospholipid transport (e.g., NIR2/PITPNM1, APOA4), protein metabolism (e.g., EIF3B), and neurodegenerative diseases/oxidative phosphorylation (e.g., COX6C) (Figure 7C and Figure 7D; Table S7G). Again, some of these male-specific hits seem to show some level of genotype difference also in females (i.e., NIR2/PITPNM1 and APOA4; Figure 7D). Globally, NLGN2 absence triggers considerable changes to the motor cortex proteome of both females and males, with a relatively low proportion of proteins commonly modified in both sexes (∼14.5%), which reminds effects on the transcriptome.

With the goal to identify some core functional elements dysregulated in *Nlgn2* KO female and male mice, common hits between the 389 DEGs (FDR < 0.05 in females and/or males) and the 716 DEPs (p < 0.05 in females and/or males) were extracted. This resulted in the identification of 30 hits having both gene expression and protein level modified in the absence of NLGN2 among which elements linked to cytoskeleton and structural remodeling (e.g., *Camkv*/CAMKV, *Ank1*/Ankyrin-1, *Acan*/Aggrecan), the MAPK pathway (e.g., *Map3k5*/MAP3K5), neuroinflammation (e.g., *Fbxl16*/FBXL16, *Lgals1*/LGALS1), and neurotransmission and neuroplasticity (e.g., *Lrrtm1*/LRRTM1, *Syt2*/SYT2, *Sv2c*/SV2C, *Gad1*/GAD1 or GAD67) (Figure 8A to 8E). Not surprisingly, some of these elements show genotype-driven changes that were larger in amplitude or only significant in one sex when considering mRNA and/or protein (e.g., *Synpo*/SYNPO, *Mtus2*/TIP150, *Scn1a*/SCN1A). It is also interesting to note that some of these hits show effects of genotype that are different when considering gene expression versus protein especially in males (e.g., *Plcxd2*/PLCXD2, *Nenf*/Neudesin, *Map3k5*/MAP3K5). For instance, Neudesin (a neurotrophin acting via the MAPK signaling pathway) shows a lower gene expression but a higher protein level in the cerebral cortex of *Nlgn2* KO mice (Figure 8C). When examining targets of interest not part of the core 30 hits but associated to cell adhesion and inhibitory neurotransmission, *Nlgn3*/NLGN3 was also found to be changed by genotype in opposite directions when considering gene expression and protein of males (Figure 8F). In general, for most of these hypothesis-driven targets, the effect of genotype appeared to reach (or approach) statistical significance only for protein level, and mutant mice showed lower level than WT mice (e.g., MDGA2, KCC2, *Gabra3* or GABAARα3). Of note is also that protein level was found to be lower in males compared to females for Aggrecan, TIP150, LGI2, GAD1/GAD67 and IGSF21. Overall, the complete absence of NLGN2 in mice considerably impacts the motor cortex transcriptome and proteome in a manner particularly affecting essential components of excitatory and inhibitory synapses, of specific signaling cascades (MAPK/ERK, FOXO, calcium), of elements controlling neuronal excitability and cellular metabolic pathways.

**Figure 8.**
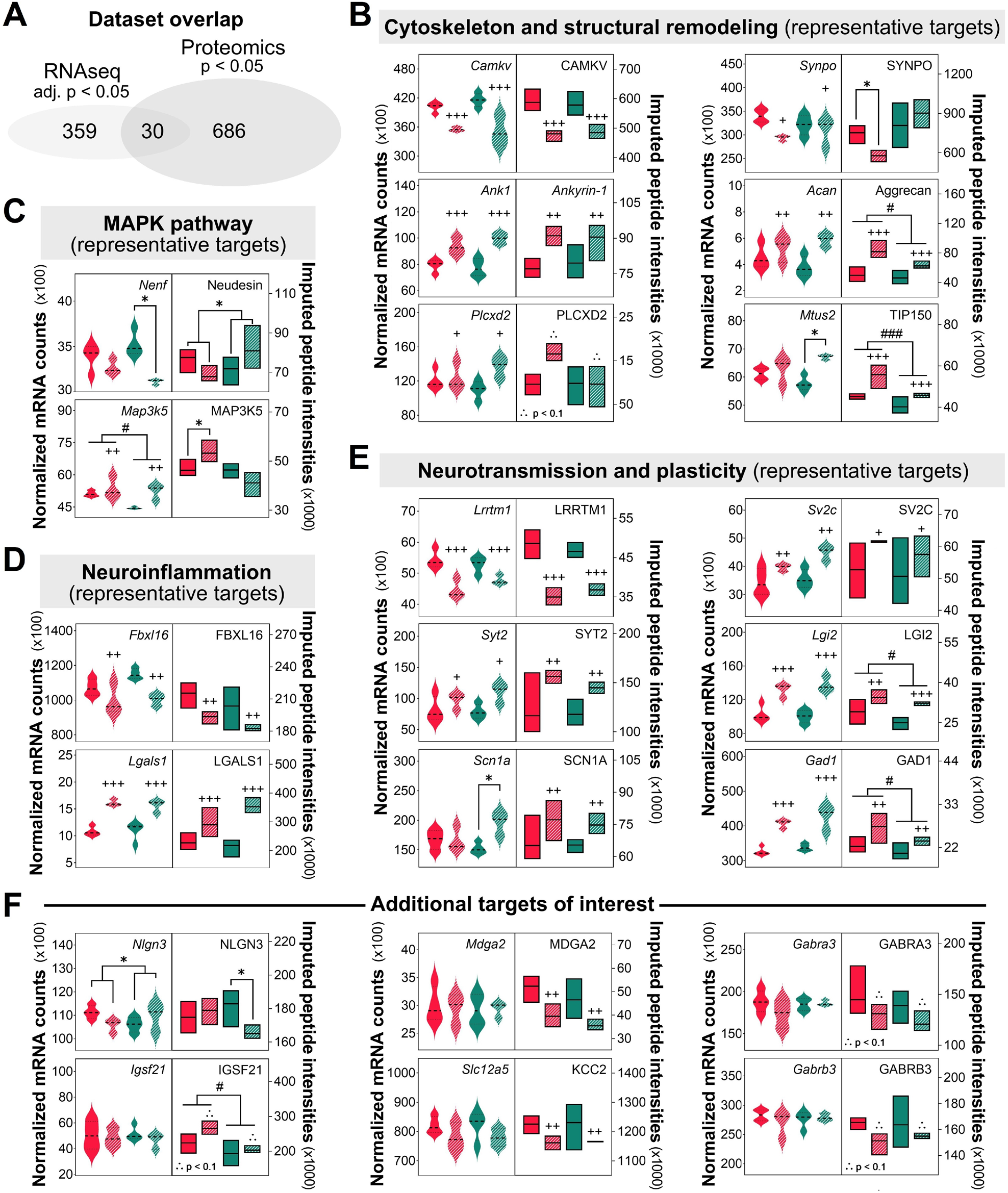
Overlap between motor cortex differentially expressed genes (DEGs) and differentially expressed proteins (DEPs) found when comparing *Nlgn2* KO female and male mice to WT littermates. **(A)** Venn diagram showing the overlap between DEGs (FDR < 0.05) and DEPs (p < 0.05) when pooling together DEGs and DEPs found in females and males. The 30 overlapping DEGs/DEPs have been used to select representative targets for panels B to E. **(B)** Expression values for selected targets related to the cytoskeleton and structural remodeling. **(C)** Expression values for selected targets related to the MAPK pathway. **(D)** Expression values for selected targets related to neuroinflammation. **(E)** Expression values for selected targets related to neurotransmission and plasticity. **(F)** Expression values for additional targets of interest defined according to previous literature (hypothesis-driven choices). Females are indicated by red/pink and males by green, with solid colors representing WT mice and hatched ones KO mice. Differences between KO and WT (females and males combined) are represented by +: p < 0.05, ++: p < 0.01 and +++: p < 0.001, and differences between females and males (genotypes combined) are represented by #: p < 0.05 and ###: p < 0.001. Differences between WT and KO males or between WT and KO females are represented by *: p < 0.05. For DEGs and DEPs, respectively, WT females n = 4 and 4, KO females n = 3 and 4, WT males n = 4 and 5, KO males n = 3 and 3.

## 4 Discussion

This study exposes the extensive changes in wake/sleep architecture, rhythmic and arrhythmic activities of the ECoG driven by the absence of NLGN2 in female mice, and unveils how these alterations compare to changes found in male mice lacking the same protein as well as the associated omic landscapes. More precisely, KO animals of both sexes spent less time in SWS and PS, and showed more fragmentation of states during the light period than WT animals. These changes in wake/sleep architecture were generally less pronounced for KO females than KO males. Moreover, mice lacking NLGN2 displayed widespread alterations in ECoG spectral power during all vigilance states, with KO females expressing larger changes during PS. When considering the dynamics of SWS delta activity and of SW density/properties, KO animals of the two sexes were almost indistinguishable, notably expressing a lower amplitude of 24-hour variations, an attenuated response to SD, and a higher density, amplitude and slope of SWS SW in comparison to WT. Modifications of multifractal metrics for all states in animals lacking NLGN2 were also very similar between sexes, and notably pointed to more self-similarity across ECoG frequencies in comparison to WT. Altogether, the data support a major effect of the *Nlgn2* mutation in both sexes, accompanied by subtle differences between males and females in the effect of genotype. The findings (obtained here with C57BL/6 mice) are very similar to those reported previously in male mice carrying the *Nlgn2* mutation on a mixed genetic background (Leduc et al., 2024; Leduc et al., under review; Seok et al., 2018), emphasizing the important role of NLGN2 in sleep regulation. Importantly, the results also reveal the sizeable impact of NLGN2 absence on genome-wide gene expression, particularly in males, and on the proteome of the cerebral cortex, providing unique indications on mechanisms underlying ECoG phenotypes.

Substantial alterations in wake/sleep architecture and in the transcriptome were found in *Nlgn2* KO animals of both sexes in the current study (e.g., less time spent and shorter episodes for SWS and PS), with some changes more important when considering males under undisturbed conditions (i.e., decreased PS and higher number of wake and SWS episodes during the light period, higher number of DEGs). These observations are reminiscent of manipulations negatively impacting NLGN2 function that were reported to trigger anxiety-like behaviors in mice of both sexes (Blundell et al., 2009; Pandey et al., 2025) or predominantly in males (Chen et al., 2017). The decreased time asleep under genetic deletion of NLGN2 could be driven by a reduced influence of sleep-promoting brain regions such as the ventrolateral preoptic area that has inhibitory (GABAergic-) projections to wake-promoting regions like orexinergic neurons of the lateral hypothalamus and histaminergic neurons of the tuberomammillary nucleus (Brown et al., 2012; Prokofeva et al., 2023; Sherin et al., 1998). Given that NLGN2 predominantly locates at GABAergic synapses (Chih et al., 2006; Varoqueaux et al., 2004), recruits/clusters gephyrin and GABA_A_ receptors (Poulopoulos et al., 2009), and regulates inhibitory synaptic transmission (Chen et al., 2020; Groisman et al., 2023; Lech et al., 2026; Liang et al., 2015; Pandey et al., 2025; Troyano-Rodriguez et al., 2019), NLGN2 absence might indeed generate a reduced inhibition of wake-promoting regions and thus an increased time spent awake, as observed under chemical lesion of the ventrolateral preoptic area (Eikermann et al., 2011). Changes in the functioning of projections to orexinergic neurons could be of particular relevance to explain some of the larger effects found in mutant males since larger effects in males in comparison to females were reported following perturbation of orexinergic transmission in mice (Dawson et al., 2023) [but not always (Durairaja & Fendt, 2021)] and more orexin-immunoreactive cells are found in C57BL/6 males than in females (Brownell & Conti, 2010). Given that orexin neurons project to the motor cortex (Du et al., 2025), sex-specific dysregulation in KO mice could even contribute to sex-specific changes in the transcriptome of the motor cortex. In parallel, a decreased inhibitory neurotransmission induced by NLGN2 absence can also increase time spent awake via negatively impacting outputs of parafacial zone GABAergic neurons promoting SWS (Anaclet et al., 2014). Alterations in these circuits are potentially also contributing to observations related to the consolidation/fragmentation of states in *Nlgn2* KO mice since, for instance, orexinergic transmission was reported to regulate transitions between states and the maintenance of arousal (Adamantidis et al., 2007; Chowdhury et al., 2019; Li et al., 2022). Future studies should investigate the role of NLGN2 in these circuits in wake/sleep regulation using cell type-specific manipulations of NLGN2 in female and male mice.

Differences in time spent in states and wake/sleep consolidation/fragmentation in the absence of NLGN2 were robust to the effect of a homeostatic sleep challenge. Indeed, more time spent awake and less time spent in SWS and PS were also found during the dark period of the recovery 24 hours in KO mice together with shorter episodes of sleep states. Under these challenged conditions, females and males did not significantly differ for the effect of the mutation further supporting a robust role of NLGN2 in shaping wake/sleep states. SWS episode duration was previously proposed as a marker of homeostatic sleep regulation that associates with the activity of GABAergic neurons of the cerebral cortex expressing nitric oxide synthase (Morairty et al., 2013). This study also highlighted a potential contribution of this neuronal population in coupling SWS episode duration to SWS intensity after SD as measured by delta power rebound, showing that animals lacking neuronal nitric oxide synthase express shorter individual SWS episodes and a reduced delta power response to SD (Morairty et al., 2013). These observations remind that of female and male mice lacking NLGN2, which can suggest a contribution of NLGN2 in the output signals of this specific GABAergic neuronal population of the cerebral cortex, one of the few neuronal populations of the brain activated during sleep (Gerashchenko et al., 2008). This hypothesis could be supported by the current findings of modifications in the level of GABAergic neurotransmission-related proteins in the motor cortex of KO mice (e.g., *Gabra2*/GABAARα2, *Gad1*/GAD67, KCC2). Interestingly, alterations in nitric oxide producing neurons was previously found in animals carrying a mutation of NLGN3 (Sharna et al., 2020), a relationship that is worth exploring also for NLGN2 in the cerebral cortex of both female and male mice, especially given the indication that the function of NLGN3 at inhibitory synapses depends on NLGN2 (Nguyen et al., 2016), and the current finding of lower level of NLGN3 in the motor cortex of *Nlgn2* KO male mice.

The absence of NLGN2 was found to drive massive changes in absolute spectral power across a wide range of frequencies in all three vigilance states almost independently of sex. These alterations may reflect changes in the cytoarchitecture or circuit wiring of the cerebral cortex impacting synchronized firing rhythms across states given the elevated power in KO mice for most frequencies in all states. This suggestion, in particular regarding cytoarchitecture, could be supported by the current finding of higher protein level of key components of the extracellular matrix in the motor cortex of mutant animals of both sexes, for instance the perineuronal net-associated proteins Ankyrin-1 (or Ankyrin-R) and Aggrecan (Frischknecht and Gundelfinger, 2012; Stevens et al., 2021). Interestingly and as reported before (Leduc et al., 2024), the shape of the spectral signature of PS (and also of wakefulness) was modified in *Nlgn2* mutant females and males, showing a lower peak frequency in the theta range. The hippocampus is a main contributor to theta activity during wakefulness and PS (Boyce et al., 2016; Fournier et al., 2020), and is tightly connected to the activity of cortical neurons (Fournier et al., 2020). It is thus speculated that hippocampal circuit alterations under NLGN2 dysregulation in mice of both sexes are contributing to changes in the spectral signature of vigilance states. Altered hippocampal function driven by NLGN2 manipulation is heavily supported by previous research (e.g., Jedlicka et al., 2011; Van Zandt et al., 2019; Wang et al., 2025). In parallel, the activity of GABAergic neurons of the median raphe correlates with hippocampal theta oscillations (Huang et al., 2022; Jelitai et al., 2021), likely modulating PS theta activity via locally shaping the activity of serotonergic and glutamatergic median raphe neurons (Huang et al., 2022), which project to the hippocampus (Fortin-Houde et al., 2023; Xu et al., 2021). Thus, NLGN2 absence could disrupt GABAergic transmission within the median raphe and, consequently, influence hippocampal theta rhythm during wakefulness and PS, which could be investigated by targeting NLGN2 downregulation to specific neuronal populations of the median raphe.

In response to elevated sleep pressure, *Nlgn*2 KO animals showed, independently of sex, a major attenuation of the increased delta power during early recovery sleep. Wakefulness dominated by theta activity was previously found to contribute to the level of delta power in subsequent sleep (Vassalli & Franken, 2017). As such, the major diminution in KO mice of the increase in theta (and alpha) activity during SD found in WT littermates could contribute to the blunted delta power response during recovery sleep. The blunted delta power response to SD in mutants may originate from some level of saturation in firing synchrony in the delta range preventing further increase under elevated homeostatic sleep pressure since the absence of NLGN2 is accompanied by a higher absolute delta power and higher density, amplitude and steeper slope of SW during SWS. More SW, larger amplitude SW, steeper SW slope and shorter negative duration of SW are compatible with the higher general excitability of neuronal circuits under the absence of NLGN2 that has been reported in previous research (Jedlicka et al., 2011; Troyano-Rodriguez et al., 2019). Importantly, our findings suggest that such a hyperexcitability is equally found in males and females under NLGN2 dysregulation, a comparison not often depicted in available research, and that it could be driven by a higher level of voltage-gated sodium channels (e.g., SCN1A) or of proteins involved in synaptic vesicle release (e.g., SYT2) in the cerebral cortex. A higher excitability of neuronal networks in the absence of NLGN2 is also likely to contribute to the altered relationship of the activity across frequencies of the interrogated spectrum indexed by the higher multifractal metric *Hm* found in KO mice. Mutant mice indeed showed a higher self-similarity across time scales (long-range dependency) in all three vigilance states, under both undisturbed and challenged conditions. Studies have suggested that the aperiodic component of synchronized brain activity might reflect the ratio between excitation and inhibition of neuronal networks (Brake et al., 2024; Gao et al., 2017). Such a ratio will be higher in the case of higher excitability and can originate from increased excitation or decreased inhibition or both. In the absence of NLGN2, inhibitory synaptic transmission is impaired (Jedlicka et al., 2011; Poulopoulos et al., 2009); and if the inhibition of interneurons projecting to pyramidal excitatory neurons is consequently decreased, it can also lead to a higher excitation. In fact, the presented gene ontology analyses of the motor cortex transcriptome and proteome seem to converge in supporting substantial changes to both inhibitory and excitatory synapses in *Nlgn2* KO mice. In the context of SWS only, the dispersion of Hurst exponents around *Hm* was higher in KO females and males, which can point to a larger number of local fractal dynamics in the between-frequency organization of the ECoG during that state. The brain aperiodic activity has been associated with the quality and speed of information processing in humans (Euler et al., 2024; Krystecka et al., 2024). Accordingly, the identification of modifications in aperiodic brain activity in an animal model of neurodevelopmental disorder is of interest to help understanding the molecular determinants contributing to an altered computational treatment in an unhealthy brain (see also Leduc et al., under review).

A crucial contribution of the present work concerns the exposition of omic signatures of the NLGN2-deficient motor cortex, which, to our knowledge, has never been reported for any brain region in this widely used animal model. The findings thus provide much-required reference datasets for the field of neurodevelopment, especially given the careful consideration of both biological sexes. The effect of the *Nlgn2* KO genotype on the transcriptome appears considerably larger than that of mutations in other cell/synaptic adhesion proteins such as *Nlgn1* and *EphA4* (Freyburger et al., 2016; Massart et al., 2014). This can be driven by a more important role of NLGN2 in shaping cellular and synaptic function, as also supported by the major changes in wake and sleep phenotypes, or by a more sensitive methodological approach used in the current study (i.e., RNA sequencing versus microarray, motor cortex versus full forebrain). It is interesting to note that many of the molecular and cellular functions impacted by NLGN2 absence in the motor cortex transcriptome and proteome (e.g., inhibitory and excitatory neurotransmission, protein phosphorylation, neuronal plasticity), are comprised in the core genome-wide pathways associated to the regulation of sleep (Mongrain et al., 2025). Therefore, a considerable number of changes found in *Nlgn2* KO mice are likely to contribute to the altered ECoG features. Among pathways that could orchestrate cellular and synaptic changes driving diverse ECoG modifications, MAPK/ERK signaling could represent a key hub. It was indeed previously linked to both ECoG activity and neurodevelopment disorders (Ballester Roig et al., 2021; Iroegbu et al., 2021), and shows dysregulation in the absence of NLGN2 when considering the transcriptome and proteome. In fact, the current alignment of the transcriptome and proteome presents a unique strategy to identify alterations at different levels of functional genomics. Moreover, it was useful to identify sex-specific omic signatures of *Nlgn2* KO mice, which seems pertinent to MAPK/ERK signaling but also to other relevant elements. Indeed, NLGN2 absence was found to modulate a large ensemble of hits tightly linked to neurodevelopmental disorders including some lower only in mutant females (e.g., SYNGAP1, SHANK3) and other lower in mutant females and males (e.g., MDGA2, CACNA1E, KCC2). Many of these targets were previously linked to alterations in ECoG oscillations (e.g., SYNGAP1 in Berryer et al., 2016), which unveils a path by which larger ECoG disturbances could be driven in mutant females.

A first potential limitation of the current study is that the phase of the estrous cycle of female mice was not monitored. Even if the effect of the estrous cycle on wake/sleep architecture and ECoG activity was reported to be relatively minor in C57BL/6 mice (Dib et al., 2021; Koehl et al., 2003), the effects of the *Nlgn2* mutation could vary according to the phase of the estrous cycle, which consideration could have revealed additional genotype differences in females. A second limitation could reside in the use of a constitutive KO animal that can give rise to compensatory mechanisms occurring notably during development. It was shown in a previous and the current study that *Nlgn1/2/3* triple KO mice and *Nlgn2* KO mice display alterations in many other synaptic proteins, such as NMDA receptors, synaptobrevin 2, synaptotagmin 1 and 2, synaptophysin 1, VGluT1 and KCC2 (Varoqueaux et al., 2006). These could contribute to wake/sleep phenotypes (as discussed above for some hits) in a manner that could differ from the role of NLGN2 in normally developed animals. The constitutive mutant also involves abnormal functioning of multiple brain regions and circuits likely contributing differently to the regulation of wakefulness and sleep. Nevertheless, this transgenic mouse presents value in the context of neurodevelopmental and neuropsychiatric disorders that (often) associate with genetic modifications impacting both nervous system development and function. A last limitation of the current work concerns the interrogation of the full tissue transcriptome and proteome preventing the understanding of the effects on different cell types. The presented omic datasets can already suggest a particular impact on different neuronal subtypes, notably parvalbumin (PVALB) interneurons in male KO mice, rather than for glial cells given the absence of significant genotype effects for many glial markers (e.g., *Gfap* and GFAP, *Aldh1l* and ALDH1L, S100, *Olig2*). Nevertheless, the use of single-cell methodologies in the future will be required to clarify the extent of the *Nlgn2* KO-driven effects on the functional genomic outputs of different brain cells.

To conclude, *Nlgn2* KO mice of both sexes show a wide range of alterations in wake/sleep architecture, in ECoG spectral and aperiodic activities, as well as in omic landscapes. The phenotypes described here are consistent with previous reports in a similar KO animal (Leduc et al., 2024; Leduc et al., under review; Seok et al., 2018), and impacts of NLGN2 absence on wake/sleep phenotypes in females and on the transcriptome and proteome of the motor cortex are here newly reported. Given that NLGN2 has been linked to neurodevelopmental and neuropsychiatric conditions (e.g., ASD showing a higher prevalence in males; Loomes et al., 2017), which are known to associate with sleep disturbances in a manner that depends on sex (Bricout et al., 2024; Estes et al., 2023), the current study highlights a molecular route associated to a diversity of modifications in brain gene and protein expression by which sex differences can occur when considering disturbances in wakefulness and sleep found in neurodevelopmental/neuropsychiatric diseases. A recent study has underscored the importance of considering separately females and males to detect sleep-related alterations in preclinical research (Mannino et al., 2024). The present data aligns with this in highlighting the requirement to use both biological sexes to evaluate the impact of genetic manipulations in models of human neuropathologies.

## Acknowledgements

The authors are thankful to Chloé Provost for help regarding surgeries and post-surgery care, the staff of the Animal facility and Molecular pathology platforms of the *Centre de recherche du Centre hospitalier de l’Université de Montréal* for animal importation, animal care or RNA quality control, and the staff of Genome Québec (in particular Sarah Piché-Choquette) and of the Proteomic Platform of the Research Institute of the McGill University Health Center for omic quantifications. The authors are also grateful to Clément Bourguignon for help with multifractal analyses and to Marcos G. Frank for feedback on the interpretation of the results.

## Authors’ contributions

NL analyzed and interpreted the wake/sleep data, and wrote the first draft of the manuscript. TL, JDG and VM performed the experiments. TL and VM contributed to the optimization of slow wave detection. TL and JML contributed to multifractal analysis. TL analyzed the transcriptomic and proteomic data. VM designed the experiment, interpreted the data, provided funding, and revised the manuscript. All authors have approved the final manuscript.

## Availability of data and materials

All data will be made available upon reasonable request. Transcriptomic data will be accessible via Gene Expression Omnibus upon publication of the manuscript (accession #GSE335476).

## Consent for publication

Not applicable

## Data availability statement

All data will be made available upon reasonable request.

## Ethics approval statement

Experimental procedures were conducted according to the guidelines of the Canadian Council on Animal Care and were approved by the *Comité d’éthique de l’expérimentation animale du Centre intégré universitaire de santé et services sociaux du Nord-de-l’Île-de-Montréal* or by the *Comité institutionnel de protection des animaux* of the *CRCHUM*.

