## Supplementary Figures for "Transcriptome and proteome associated to the sex-dependent effects of Neuroligin-2 absence on wakefulness and sleep features"

Revised manuscript submitted to: *bioRxiv*

### Supplementary figures 1 and 2

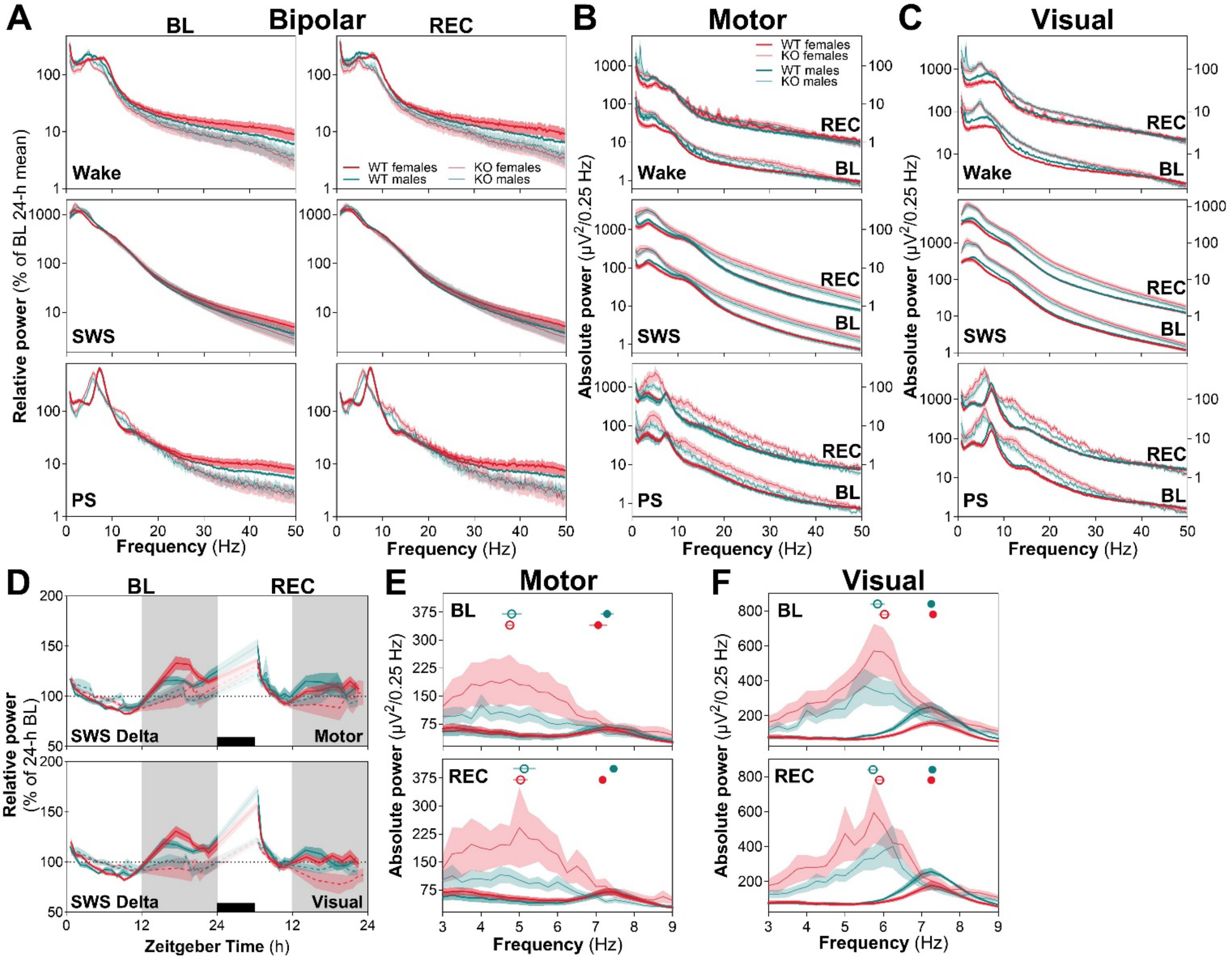


**Supplementary figure 1** - Relative power spectra computed from the bipolar signal and absolute power spectra for the motor and visual cortex signals shown for wakefulness, slow wave sleep (SWS), and paradoxical sleep (PS) during baseline (BL) and recovery (REC), and SWS delta activity 24-hour dynamics for the motor and visual cortex in *Nlgn2* KO females and males and WT littermates. **(A)** Relative spectral power (log scale) between 0.75 and 50 Hz (expressed as a percentage of BL 24-hour mean). WT females n = 11, WT males n = 9, KO females n = 10, KO males n = 10. **(B)** Absolute spectral power (log scale) between 0.75 and 50 Hz for the signal of the electrode placed above the motor cortex. WT females n = 10, WT males n = 7, KO females n = 10, KO males n = 10 (also for panels D [top] and E). **(C)** Absolute spectral power (log scale) between 0.75 and 50 Hz for the signal of the electrode placed above the visual cortex. WT females n = 11, WT males n = 8, KO females n = 10, KO males n = 10 (also for panels D [bottom] and F). **(D)** Time courses of relative SWS delta (1-4 Hz) activity for the motor and visual signals. Gray backgrounds indicate the 12-hour dark periods and black rectangles the 6-hour sleep deprivation (SD). **(E)** Zoom on the theta frequency range of the PS absolute ECoG power spectrum of the motor signal. **(F)** Zoom on the theta frequency range of the PS absolute ECoG power spectrum of the visual signal. For panels E and F, PS peak frequency is displayed as open (KO mice) and closed (WT mice) circles and red and turquoise represent, respectively, female and male animals.

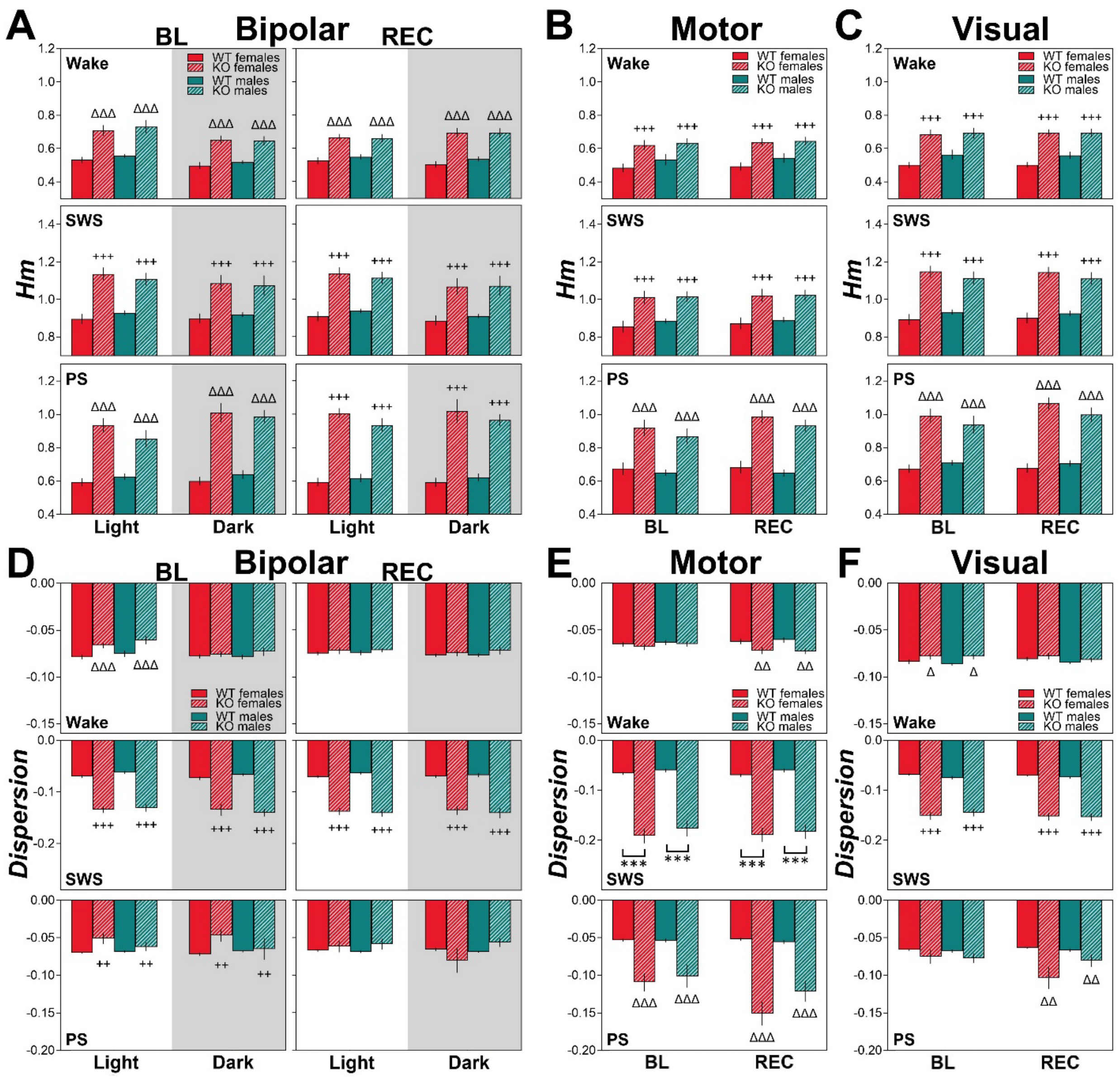


**Supplementary figure 2** - Most prevalent Hurst exponent (*Hm*) and *Dispersion* of exponents around *Hm* computed for wakefulness, slow wave sleep (SWS), and paradoxical sleep (PS) during baseline (BL) and recovery (REC) in *Nlgn2* KO mice and WT littermates when considering the light and dark periods separately for the bipolar signal or the full 24-hour days for the signals of the motor or the visual cortex. **(A)** *Hm* averaged for the 12-hour light and 12-hour dark periods during BL and REC. WT females n = 11, WT males n = 9, KO females n = 10, KO males n = 10. **(B)** *Hm* for the motor cortex signal averaged over 24 hours for BL and REC. WT females n = 10, WT males n = 7, KO females n = 10, KO males n = 10. **(C)** *Hm* for the visual cortex signal averaged over 24 hours for BL and REC. WT females n = 11, WT males n = 8, KO females n = 10, KO males n = 10. **(D)** *Dispersion* averaged for the 12-hour light and 12-hour dark periods during BL and REC. WT females n = 11 (except n = 10 for wake), WT males n = 8 (except n = 9 for SWS in REC), KO females n = 10, KO males n = 10. **(E)** *Dispersion* for the motor cortex signal averaged over 24 hours for BL and REC. WT females n = 10 (except n = 9 for wake), WT males n = 7 (except n = 6 for wake and PS), KO females n = 10, KO males n = 10. **(F)** *Dispersion* for the visual cortex signal averaged over 24 hours for BL and REC. WT females n = 11 (except n = 10 for wake), WT males n = 8 (except n = 7 for wake and PS), KO females n = 10, KO males n = 10. Differences between WT and KO animals encompassing the full 24-hour of BL or REC conditions (panels A and D) or the full 48-hour of BL and REC (panels B, C, E, and F) are represented by ++: $p < 0.01$ and +++: $p < 0.001$; differences between WT and KO animals that are shaped by the light/dark period (panels A and D) or by BL/REC condition (panels B, C, E, and F) are represented by Δ: $p < 0.05$, ΔΔ: $p < 0.01$, and ΔΔΔ: $p < 0.001$; differences between WT and KO males or WT and KO females are represented by ***: $p < 0.001$. Gray backgrounds indicate the 12-hour dark periods.
